# Synaptic proteins that aggregate and degrade slower with aging accumulate in microglia

**DOI:** 10.1101/2025.05.20.654652

**Authors:** Ian H. Guldner, Viktoria P. Wagner, Patricia Moran-Losada, Sophia M. Shi, Kelly Chen, Barbara T Meese, Hamilton Oh, Yann Le Guen, Nannan Lu, Pui Shuen Wong, Ning-Sum To, Dylan Garceau, Zimin Guo, Jian Luo, Michael Sasner, Andreas Keller, Andrew C. Yang, Tom Cheung, Tony Wyss-Coray

**Author notes:** These authors contributed equally (Andreas Keller, Andrew C. Yang, Tom Cheung).

## Abstract

Neurodegenerative diseases affect 1 in 12 people globally and remain incurable. Central to their pathogenesis is a loss of neuronal protein maintenance and the accumulation of protein aggregates with aging^1,2^. We engineered bioorthogonal tools^3^ which allowed us to tag the nascent neuronal proteome and study its turnover with aging, its propensity to aggregate, and its interaction with microglia. We discovered neuronal proteins degraded on average twice as slowly between 4- and 24-month-old mice with individual protein stability differing between brain regions. Further, we describe the aged neuronal ‘aggregome’ encompassing 574 proteins, nearly 30% of which showed reduced degradation. The aggregome includes well-known proteins linked to disease as well as a trove of proteins previously not associated with neurodegeneration. Unexpectedly, we found 274 neuronal proteins accumulated in microglia with 65% also displaying reduced degradation and/or aggregation with age. Among these proteins, synaptic proteins were highly enriched, suggesting a cascade of events emanating from impaired synaptic protein turnover and aggregation to the disposal of these proteins, possibly by the engulfment of synapses by microglia. These findings reveal the dramatic loss of neuronal proteome maintenance with aging which could be causal for age-related synapse loss and cognitive decline.

## Main Text

Aging is accompanied by a loss of proteostasis, the maintenance of a balanced and functional proteome, a major contributor to organismal deterioration associated with aging^1,4^. All aspects of proteostasis have been observed to be disrupted with aging, including the ability of the cell to maintain proper protein synthesis and degradation rates, transport of proteins to their proper destinations, and prevent the accumulation of misfolded and aggregated proteins^1,4^. The loss of proteostasis in the brain is a major contributor to age-associated vulnerability to reduced cognitive and motor abilities and neurodegenerative diseases^2^. Indeed, compromising broad proteostasis pathways in experimental models has demonstrated such manipulations can induce dementia-like phenotypes^2,5,6^. Understanding the dynamics of proteostasis in a neuron-specific manner could help define mechanisms or individual proteins that could be exploited for therapeutic purposes. However, despite the emergence and application of some tools to study cellular proteomes^3,7–16^, such research has been hindered by a lack of robust models to examine protein dynamics in a cell-specific manner in higher organisms. Here, we develop new *in vivo* models that enable robust bioorthogonal tagging of nascent proteomes in a cell-specific manner by expression of mutant aminoacyl-tRNA synthetases (aaRSs). We leveraged these models to study key features of neuronal proteostasis dynamics with aging, including protein degradation, protein aggregation, and neuronal-to-microglia protein transfer, culminating in unparalleled insight into the decline of neuronal proteostasis with age and the identification of a microglia-mediated mechanism of maintaining neuronal proteostasis.

### *In Vivo* Cell-Specific Nascent Proteome Labeling by New BONCAT Models

Expanding upon our earlier *in vitro* studies^3^, we generated two new bioorthogonal non-canonical amino acid tagging (BONCAT) knock-in mouse lines with cassettes expressing mutant aaRSs, flox-stop-flox-eGFP-p2a-PheRS^T413G^ and flox-stop-flox-eGFP-p2a-TyrRS^Y43G^ (hereafter, PheRS* and TyrRS*, respectively) (**Fig. 1a**). We first aimed to compare the protein tagging efficacy of our models to that of the current standard BONCAT transgenic mouse line based on the expression of a mutant methionine aaRS (hereafter, MetRS*)^7,8^. Each of the three BONCAT lines were crossed to a Camk2a-Cre driver and administered its respective azido-modified amino acid (AzAA) to evaluate nascent proteome labeling in excitatory neurons^17^ (**Fig. 1a, 1b**). By in-gel fluorescence, the Camk2a-Cre;PheRS* model showed a high fluorescence signal over its respective background control while the Camk2a-Cre;TyrRS* and Camk2a-Cre;MetRS* did not show an appreciable difference in fluorescence relative to their respective background controls (**Fig. 1c**). Treating Camk2a-Cre;MetRS* mice with AzAA-infused water while fed a low-methionine chow as originally reported did not appreciably improve the signal-to-noise ratio (**Extended Data Fig. 1a**)^8^. These results were mirrored by *in situ* tissue staining for azide-modified proteins (**Fig. 1d**). The fluorescence signal observed in Camk2a-Cre;PheRS* brain sections co-localized to GFP+ cells possessing neuronal morphology (**Extended Data Fig. 1b**) with the expected spatial distribution of Camk2a+ neurons based on *in situ* hybridization data (**Fig. 1d, Extended Data Fig. 1c**). Lastly, we tested proteome labeling of the three BONCAT models by performing liquid chromatography-mass spectrometry (LC-MS) on BONCAT-labeled proteins enriched by bead-based pulldown (**Extended Data Fig. 1d**). We found the different models separated well by PCA (**Fig. 1e**). We detected 3,787 proteins in Camk2a-Cre;PheRS*, 2,320 proteins in Camk2a-Cre;MetRS*, and 4 proteins in Camk2a-Cre;TyrRS* (**Fig. 1f**). Of note, 1,959 proteins were commonly detected between Camk2a-Cre;PheRS* and Camk2a-Cre;MetRS*, with 1,828 proteins being uniquely detected in Camk2a-Cre;PheRS* and only 361 proteins uniquely detected in Camk2a-Cre;MetRS* (**Fig. 1f**). The statistical confidence of protein detection (**Fig. 1g, Extended Data Fig. 1e**) and signal-to-noise ratio (**Fig. 1h, Extended Data Fig. 1f**) was significantly higher in Camk2a-Cre;PheRS* compared to the other two BONCAT lines. The robust, maximal labeling in the Camk2a-Cre;PheRS* (**Extended Data Fig. 1g, 1h**) did not induce HSP90 expression (**Extended Data Fig. 1i**) or microgliosis (**Extended Data Fig. 1j**), suggesting azide-modified residues do not induce proteostasis stress or an immune response, consistent with similar observations in MetRS* models^7,10^.

**Figure 1:**
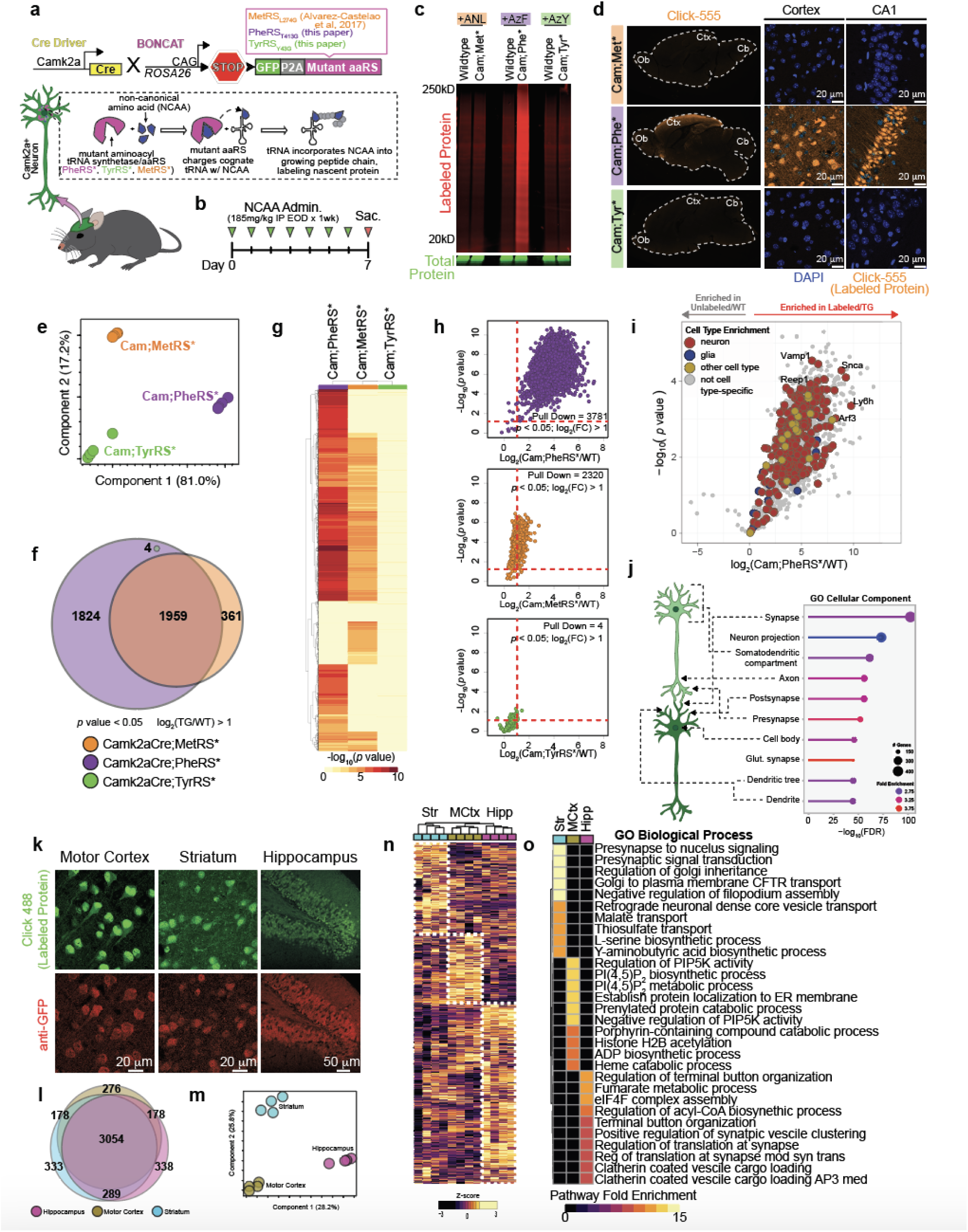
Evaluation of Nascent Proteome Labeling of Camk2a+ Excitatory Neurons by BONCAT Mouse Lines. a. Schematics of transgenes to permit mutant amino-acyl tRNA synthetase (aaRS) expression in mice to allow bioorthogonal non-canonical amino acid tagging (BONCAT) and the mechanism of nascent proteome labeling in BONCAT mice. b. Timeline of non-canonical amino acid (NCAA)/azido-amino acid (azAA) administration for nascent proteome tagging in BONCAT transgenic mice. c. In-gel fluorescence image of Alexa 647-clicked and thus BONCAT labeled proteins from total brain lysates derived from young Camk2aCre;MetRS*, Camk2aCre;PheRS*, and Camk2aCre;TyrRS* BONCAT transgenic mice and their respective wildtype background controls. Relative Alexa 647 intensities can be taken as a measure of relative labeling efficiencies. d. Fluorescence images of Alexa 555-clicked and thus BONCAT labeled proteins in brain tissue sections from young Camk2aCre;MetRS*, Camk2aCre;PheRS*, and Camk2aCre;TyrRS* BONCAT transgenic mice. e. Principal component analysis based on the abundance of BONCAT labeled proteins from young Camk2aCre;MetRS*, Camk2aCre;PheRS*, and Camk2aCre;TyrRS* BONCAT transgenic mice as determined by LC-MS. Proteins not identified in one group but identified in others were maintained for analysis and values imputed. n = 4 mice per experimental group. f. Venn Diagram comparing the number of different proteins commonly and exclusively identified by LC-MS in young Camk2aCre;MetRS*, Camk2aCre;PheRS*, and Camk2aCre;TyrRS* BONCAT transgenic mice. n = 4 mice per experimental group. Proteins used in the comparison had a log_2_ fold change > 1 over the respective background control with a *p* value < 0.05. g. Heatmap comparing -log_10_ *p* values of identified proteins in young Camk2aCre;MetRS*, Camk2aCre;PheRS*, and Camk2aCre;TyrRS* BONCAT transgenic mice. n = 4 mice per experimental group. Proteins not identified in a particular line were assigned a -log_10_ *p* value of 0. h. Volcano plots showing the relative abundance and *p* values of proteins detected by LC-MS in young Camk2aCre;MetRS*, Camk2aCre;PheRS*, and Camk2aCre;TyrRS* BONCAT transgenic mice relative to their respective wildtype background controls. n = 4 mice per experimental group. Each volcano plot can be taken as a measure of the signal-to-noise ratio between each BONCAT mouse line and the respective background control. i. Color-coded volcano plot showing the enrichment of BONCAT-labeled proteins identified by LC-MS in young Camk2a-Cre;PheRS* BONCAT mouse line relative to the respective wildtype background control. n = 4 mice per experimental group. Proteins are color-coded by cell type enrichment. j. Gene Ontology Cellular Component analysis on BONCAT labeled proteins in young Camk2a-Cre;PheRS* BONCAT mouse line. Proteins used in the analysis had a log_2_ fold change > 1 over the respective background control with a *p* value < 0.05. k. Fluorescence images of Alexa 488-clicked and thus BONCAT labeled proteins in motor cortex, striatum, and hippocampus of brain tissue sections from young Camk2a-Cre;PheRS* BONCAT mouse line. l. Venn Diagram comparing the number of different proteins commonly and exclusively identified by LC-MS in the motor cortex, hippocampus, and striatum of young Camk2a-Cre;PheRS* BONCAT mouse line. n = 4 mice per experimental group. Proteins used in the comparison had a log_2_ fold change > 1 over the respective region background control with a *p* value < 0.05. m. Principal component analysis based on the abundance of BONCAT labeled proteins from the motor cortex, striatum, and hippocampus of young Camk2a-Cre;PheRS* BONCAT mouse line as determined by LC-MS. n = 4 mice per experimental group. n. Heatmap with hierarchical clustering comparing the z-scored abundance of BONCAT-labeled proteins from the motor cortex, striatum, and hippocampus of young Camk2a-Cre;PheRS* BONCAT mouse line. Protein clusters enclosed by a white dotted line are considered region marker proteins. o. Heatmap comparing pathway fold enrichment of the top 10 Gene Ontology Biological Processes of the motor cortex, striatum, and hippocampus based on the region marker proteins shown in (n). Only pathways with an FDR < 0.05 were considered in the analysis.

The relatively poor performance of the Camk2a-Cre;TyrRS* line was unexpected as we observed the TyrRS* construct performed superiorly in labeling HEK cell proteomes *in vitro*^3^. To begin to test whether the efficacy of proteome labeling by each BONCAT line is related to the tissue examined, we performed similar experiments as above but crossed each BONCAT line to a CMV-Cre driver for ubiquitous labeling^18^. LC-MS analysis showed that depending on the tissue examined, different BONCAT lines had varying efficacy in labeling tissue proteomes (**Extended Data Fig. 1k**). For example, the CMV-Cre;TyrRS* line tagged 1102 liver proteins while the other two lines tagged less than 400 liver proteins each (**Extended Data Fig. 1k**). These data demonstrate the potential utility of all three BONCAT lines, the efficacy of which might depend on the tissue examined and the cre-driver used.

### Characterization of Camk2a+ Excitatory Neuronal Proteome with PheRS* Model

Determining the PheRS* model was optimal for labeling the Camk2a+ excitatory proteome, we further examined the neuronal proteins identified by LC-MS in this model. As expected from the Camk2a-Cre driver, many proteins (306) identified were annotated as neuronal with few (31) annotated as being specific to other cell types (**Fig. 1i**). Because many proteins are common among diverse cell types, most proteins (3463) were considered non-cell type-specific (**Fig. 1i**). Gene Ontology Cellular Component analysis revealed all major anatomical features of neurons, including the cell body, axon, dendrites, and synapses, were represented by hundreds of proteins (**Fig. 1j**). These data attest to the value of BONCAT methodology, enabling the detection of proteins from fine structural parts that would likely be sheared off during tissue dissociation and/or cell sorting, processes typically required to achieve cell-specific proteomes without cell-specific protein-tagging. In a separate cohort of labeled Camk2a-Cre;PheRS* mice, we dissected the motor cortex, striatum, and hippocampus, regions that exhibit robust labeling (**Fig. 1k**), and performed LC-MS on the enriched labeled proteins. 3054 proteins were commonly identified among all regions, but each region had 276-338 uniquely identified proteins (**Fig. 1l**). PCA analysis of the regional Camk2a+ neuronal proteomes separated all three regions (**Fig. 1m**), which was reflected by hierarchical clustering and heatmap analyses (**Fig. 1n**). Gene Ontology Biological Process enrichment showed each cluster/region was diverse in pathways, but generally represented presynaptic signaling and molecule transport for the striatum, cell membrane protein regulation and metabolic processes for the motor cortex, and multiple synaptic processes and terminal button organization for the hippocampus (**Fig. 1o**). These results point toward regional specialization of Camk2a+ neurons for particular biological processes and underscore the resolution achieved by the PheRS* model.

### Proteome-wide Reduction of Neuronal Protein Degradation with Aging Across Brain Regions

Given the fundamental role of protein turnover and aggregation in neurodegenerative diseases especially in long-lived, non-mitotic neurons^19^, we sought to determine how neuronal protein degradation differs with age in mice. BONCAT models are particularly suited to study protein turnover given their time-stamping of proteins at synthesis in a cell-specific manner, which cannot be achieved using other protein tagging methods^8,11,12,20^. To rapidly deliver the BONCAT machinery to aged mice, we developed AAV expression vectors encoding a Camk2a-driven PheRS* (**Extended Data Fig. 2a**). Mice intravenously transduced with AAV carrying the PheRS* construct showed labeling significantly higher than background controls (**Extended Data Fig. 2b**) and labeled cells possessed neuronal morphology with the expected spatial distribution of Camk2a+ cells (**Extended Data Fig. 2c**). The proteome coverage, signal-to-noise ratio, and statistical confidence of AAV:Camk2a-PheRS* mice was very similar to that of Camk2a-Cre;PheRS* transgenic mice (**Extended Data Fig. 2d, 2e**). Identified proteins were mostly neuron-specific proteins or general cellular proteins (**Extended Data Fig. 2f**) and represented all features of neuronal anatomy (**Extended Data Fig. 2g**). The AAV construct additionally labeled proteomes of aged, 21 month old mice, permitting the comparison of the aged and young neuronal proteomes. 213 proteins were differentially expressed with age, which displayed expected pathway differences between ages, such as the downregulation of neurodevelopmental pathways in aged mice (**Extended Data Fig. 2h, 2i, 2j**).

To study protein degradation changes with age, young (4 months-old), middle-aged (12 months-old), and aged (24 months-old) mice were transduced with AAV:Camk2a-PheRS* by retro-orbital injection with a pulse-chase AzF administration scheme (**Fig. 2a**). Mice were sacrificed at four time points within the two week chase period (n = 4 biological replicates per timepoint per age group) - 16 hours, 3 days, one week, and two weeks (TP1, TP2, TP3, and TP4, respectively) - and brain regions were dissected immediately upon brain extraction (**Fig. 2a**). In-gel fluorescence (**Fig. 2b**) and *in situ* tissue staining (**Fig. 2c**) both showed a dilution of tagged-protein fluorescence signal progressing through the chase period, indicative of tagged protein degradation progressing into the chase. To quantify degradation rates for individual tagged neuronal proteins across brain regions and ages, we enriched for tagged neuronal proteins as previously described, tandem-mass-tag labeled enriched peptide fractions, and ran the plexes on LC-MS (**Fig. 2a**). We obtained a kinetic degradation trajectory of the percent protein remaining over time for every protein identified, resulting in degradation profiles for hundreds of proteins for each region and each age (**Fig. 2d**). The average degradation trajectories for all proteins between regions differed, with the hippocampus and hypothalamus showing faster degradation kinetics compared to cortical regions in young mice (**Fig. 2d**). Moreover, the average degradation trajectories of aged regions relative to their respective young and middle-aged regions was broader (**Fig. 2d**), indicating slower protein degradation with aging that emerged after middle-age. These qualitative observations were further supported by quantifications of the percent protein remaining (**Fig. 2d, Extended Data. Fig 3a**).

**Figure 2:**
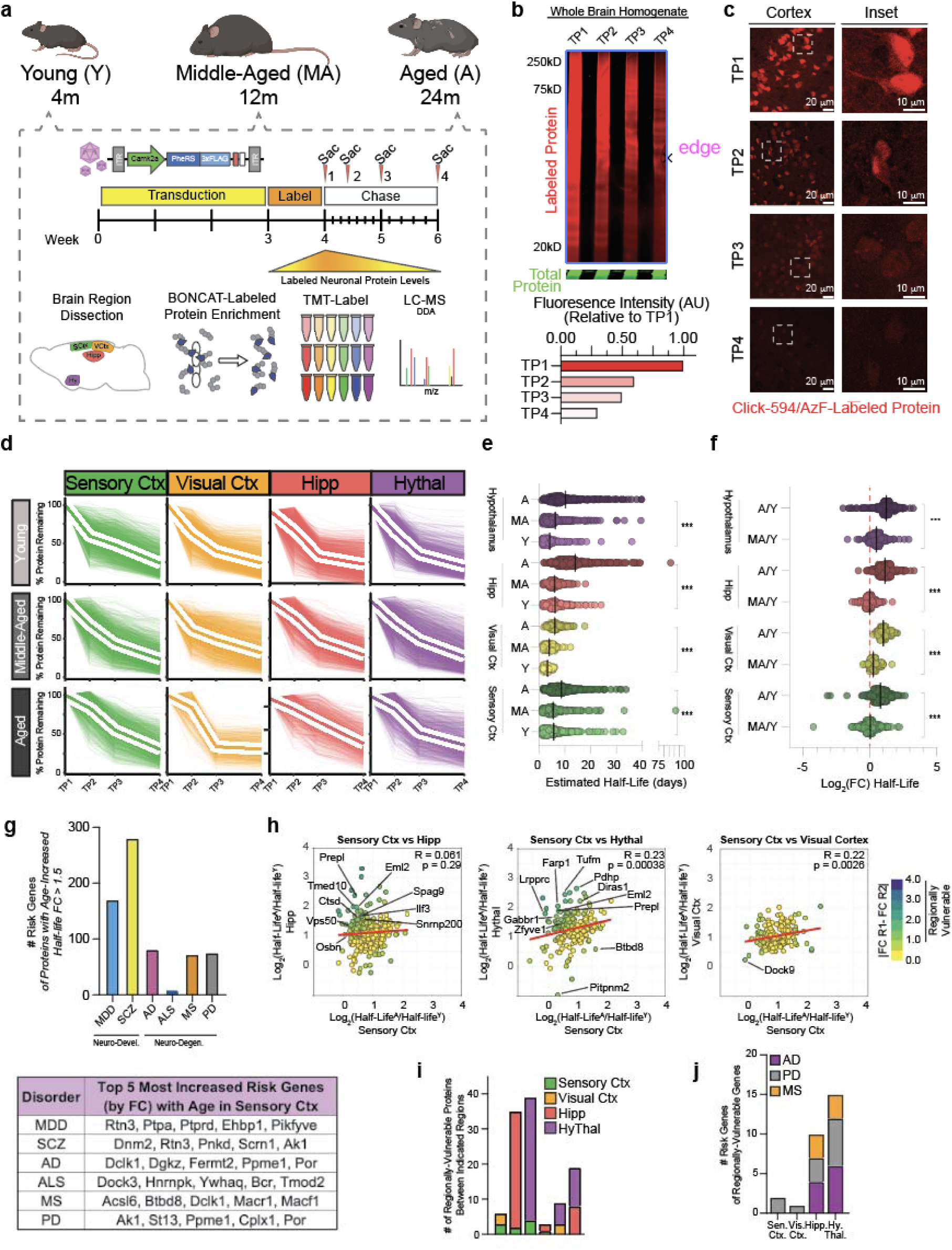
Neuronal Protein Degradation Slows with Aging and is Regionally Heterogeneous. a. Schematic of the experimental approach to study protein degradation with aging using BONCAT methodology. n = 4 mice per timepoint for each age. b. In-gel fluorescence image of Alexa 647-clicked and thus BONCAT labeled proteins of total brain lysates derived from Camk2a-Cre;PheRS* BONCAT mice at the indicated time points in the chase period following labeling of proteins with azido-phenylalanine. c. Fluorescence images of Alexa 594-clicked and thus BONCAT labeled proteins in cortex of brain tissue sections from the Camk2a-Cre;PheRS* BONCAT mice at the indicated time points in the chase period following labeling of proteins with azido-phenylalanine. d. Kinetic degradation trajectories of the percent of BONCAT-labeled protein remaining through the chase period following labeling of proteins with azido-phenylalanine in the indicated brain regions and ages. Each fine line represents one protein derived from averaging four biological replicates. The single bold line outlined in white represents the average of all proteins. Proteins were plotted irrespective of age-based or regionally-based overlap and only filtered to exclude proteins that displayed a 5% increase between any two time points. e. Plot of the estimated protein half-life in days for the indicated brain regions and ages. Each dot represents one protein. For each individual brain region, only proteins commonly identified between all ages of that region are plotted. *P* value determined by paired t-test between young and aged proteins. *** *p* < 0.0001 f. Plot of log_2_ fold change of estimated protein half-life values between the indicated brain regions and ages. Each dot represents one protein and is the same as those displayed in (f). *P* value determined by paired t-test between young and aged proteins. g. Bar plot of the number of proteins with an age-increased half-life (age vs young fold change > 1.5) that are also H-MAGMA risk genes for the indicated brain disorders. Risk genes were derived from the H-MAGMA study and considered only if the reported *p* value was < 0.05 (top). Table of the top 5 most half-life increased risk genes with age in the sensory cortex (bottom). h. Scatter plots comparing the log_2_ fold change of the estimated protein half-lives (young to aged) between proteins commonly detected between the sensory cortex versus the hippocampus (left), hypothalamus (middle), and visual cortex (right). Each dot represents one protein with the color coding representing the absolute value of the difference between log_2_ fold changes of protein half-life between the indicated regions. Proteins with an absolute value difference >1 were considered regionally vulnerable. i. Bar plot of the number of regionally-vulnerable proteins between the indicated regions. As in (h), proteins with an absolute value difference >1 were considered regionally vulnerable. j. Bar plot of the number of H-MAGMA neurodegenerative risk genes within the identified regionally-vulnerable proteins for each region analyzed. As in (g), risk genes were derived from the H-MAGMA study and considered only if the reported *p* value was < 0.05. As in (h), proteins with an absolute value difference >1 were considered regionally vulnerable.

We next estimated protein half-life by using modeling techniques established to estimate protein half-lives from protein-labeling pulse-chase proteomics data^21,22^. The estimated half-life values confirmed the qualitative observations of the kinetic curves, showing stable average half-lives from young to middle age with an increase of 2.3-8 days from middle age to aged mice, depending on the region (**Fig. 2e**). Modeled half-life values were in good correlation with direct interpolation of half-life from the non-modeled degradation trajectories (**Extended Data Fig. 3b**). The average log_2_ fold change in half-life among all regions was approximately 0.2 (∼15% increase) from young to middle-age but approximately 1.0 (100% increase/doubling) from middle-age to aged with variance between regions (**Fig. 2f**). The observation of reduced protein degradation with age is consistent with previous reports measuring protein degradation in whole mouse brain homogenates, *Drosophila melanogaster* heads, and *Caenorhabditis elegans* using SILAC methodology^23–26^. The top 10% cortical proteins with the greatest fold change increase from young to aged were enriched for proteins of the synapse (DGKZ, DCLK1, DST, SORCS2) and cell projections (CPLX1, DNM2, DST, GLUL) and for functions related to vesicle and protein transport (**Extended Data Fig. 3c, 3d**), neuronal features and pathways reported to be compromised with aging and dementias^27,28^. Intriguingly, several hundred of the proteins with >50% increase in half-life are annotated as neurodevelopmental or neurodegenerative risk genes by the Hi-C-coupled multimarker analysis of genomic annotation (H-MAGMA) study^29^ (**Fig. 2g, top**). Proteins with the most increased half-life with age that were also neurodegenerative risk genes, such as BTBD8, CPLX1, DCLK1, FERMT2, and YWHAQ (**Fig. 2g, bottom**), are proteins localized to cell junctions and the actin cytoskeleton with involvement in cell-cell junction organization and signaling, implying reduced protein degradation has repercussions for both the host cells and their signaling partners. We examined whether certain protein features, such as protein length and isoelectric point, could be ascribed to the extent of reduced degradation but found no such correlation (**Extended Data Fig. 3e**).

Next, we compared the aged-to-young half-life fold changes of proteins shared across regions, hypothesizing that proteins with differing half-life changes could contribute to regional vulnerability or resilience to aging and diseases. Comparing half-life fold changes of the sensory cortex to those of the hippocampus, hypothalamus, and visual cortex, we found no two regions were perfectly correlated (**Fig. 2h**), which was also true when making other region-wise comparisons (**Extended Data Fig. 3f**). The sensory cortex and visual cortex displayed the least extreme diversity in half-life fold change differences (n = 6 proteins, |(log_2_FC_A/YSensoryCtx_)-(log_2_FC_A/YVisualCtx_)| > 1), with more changes present when comparing the sensory cortex to either the hippocampus (n = 35 proteins) or hypothalamus (n = 39 proteins) (**Fig. 2h, 2i**). Supporting the hypothesis that these proteins could confer vulnerability or resilience, several of these proteins were risk genes as defined by the H-MAGMA study (**Fig. 2j**). The hippocampus and hypothalamus possessed 10 and 15 H-MAGMA neurodegenerative risk genes, respectively, while cortical regions had fewer than 5 each (**Fig. 2j**). Some proteins identified by this analysis, such as PREPL in the hippocampus and LRPPRC in the hypothalamus, have been experimentally demonstrated to contribute to Alzheimer’s Disease progression. However, the contribution of many other proteins, including VPS50, SPAG9, TMED10, and AMPH, to aging and neurodegeneration remains to be elucidated, but given their broad connections to protein transport and vesicle-mediated transport, likely have regulatory roles in aging and dementias.

### Coordinated Degradation of Proteins by Biological Function

Half-life values can oversimplify nuances of complex kinetic degradation trajectories such as shown in **Fig. 2d**, so we performed analyses on the kinetic degradation trajectories to potentially extract additional information from the data (**Fig. 3**). First, we clustered the degradation trajectories on a per region basis, initially clustering only profiles from young samples to serve as a baseline reference for older ages (**Fig. 3a, Extended Data Fig. 4a**). As an example, for the sensory cortex, six clusters were produced with each cluster having a visually distinguishable average profile (**Fig. 3b**) and slope values (**Fig. 3c**). The top 5 most statistically significant Gene Ontology Biological Processes terms of each cluster in the sensory cortex largely represented by one or two biological processes, such as cellular respiration for cluster 1 and synaptic vesicle cycling for cluster 5 (**Fig. 3d**), supporting the biological meaningfulness of the clustering.

**Figure 3:**
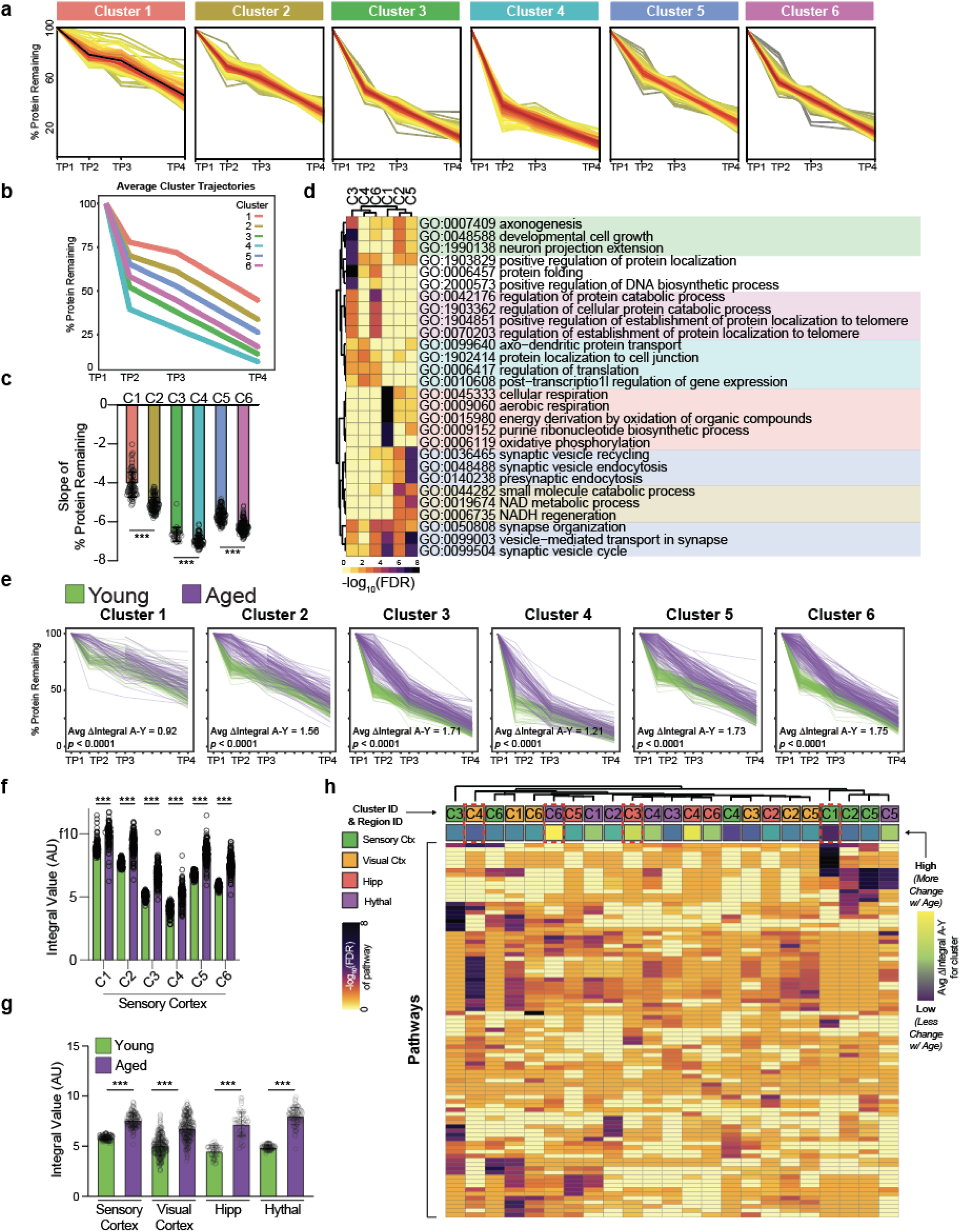
The Coordinated Degradation of Proteins by Biological Function Is Differentially Compromised with Aging. a. Kinetic degradation trajectories of the six clusters identified by unbiased clustering of all protein degradation trajectories of the young (4 months) sensory cortex. Red lines represent proteins closer to the average trajectory of the cluster while yellow lines represent those farther away from the average of the cluster. b. Plot of the average degradation trajectory for each cluster visualized in (a) from the young sensory cortex. c. Bar plot comparing the slopes of the kinetic degradation trajectories of each protein in each cluster for the young sensory cortex. Each dot represents the slope of the kinetic degradation trajectory of one protein. *P* value determined by a one-way ANOVA with significant comparisons identified by a Tukey test. d. Heatmap of the top 5 most significant Gene Ontology Biological Processes identified for each cluster in the young sensory cortex. Heatmap colors represent -log_10_ of the FDR for each pathway. e. Overlap of young (4 months) and aged (24 months) kinetic degradation trajectories of the six clusters identified in the sensory cortex with lines color-coded by age. Protein membership in the aged clusters was determined by the clustering of young samples to serve as a baseline. The average delta integral, calculated by averaging the difference of the integral values of each aged and young protein within the cluster, is reported on each plot. The *p* value was determined by a one-way ANOVA with significant comparisons identified by a Tukey test. f. Bar plot comparing the integral values of young and aged proteins within each cluster of the sensory cortex. Each dot represents the integral value for one protein within the indicated cluster. *P* value determined by a two-tailed t-test. *** *p* < 0.0001. g. Bar plot comparing the integral values of young and aged proteins on a per-region basis. *P* value determined by a two-tailed t-test. *** *p* < 0.0001. h. Heatmap of the top 5 most significant Gene Ontology Biological Processes identified for each cluster in each brain region examined. Regions and respective clusters are indicated at the top of the heatmap. Heatmap colors represent -log_10_ of the FDR for each pathway. The annotation at the top of the heatmap represents the delta integral of the indicated region/cluster.

Clusters representing distinct pathways were generally recapitulated among other brain regions (**Extended Data Fig. 4b**). Cumulatively, these results indicate that proteins within similar pathways, which likely need to be functionally coordinated with each other to execute a particular biological process, have coordinated degradation rates, an observation in-line with coordinated half-lives of individual proteins within the same protein complex^30^.

### Differential Vulnerability of Biological Pathways to Age-related Degradation Alterations

To compare how clusters change with age in the sensory cortex, we extracted aged protein profiles based on the assigned young cluster identities and subsequently overlapped the degradation profiles of aged proteins with those of young proteins on a per-cluster basis (**Fig. 3e**). Most proteins of young and aged profiles were separated (**Fig. 3e**). For the sensory cortex and all other brain regions examined, aged profiles had a larger integral value (area under the curve) than young profiles for each cluster (**Fig. 3f, 3g, Extended Data Fig. 4a**), indicative of reduced degradation rates in aged mice. When incorporating middle-aged profiles into the analysis, all regions showed an increased average integral value from middle age to aged, but only the visual cortex and hypothalamus showed increased average integral values from young to middle-aged (**Extended Data Fig. 4c**), signifying earlier degradation deficits in these regions.

We calculated the average difference of the integral values for aged and young proteins of each cluster to obtain a delta integral score for each cluster (**Fig. 3e**), a measure of the magnitude of protein degradation reduction between clusters. In the sensory cortex, Cluster 3, enriched for proteostasis and developmental processes (**Fig. 3d**), had one of the largest delta integral scores (1.71) (**Fig. 3e**) while Cluster 1, enriched for cellular respiration processes (**Fig. 3d**), had the smallest delta integral score (0.92) (**Fig. 3e**). The delta integral scores suggest that certain biological processes, such as proteostasis and developmental networks, are more vulnerable to the consequences of age-related degradation than other processes, such as cellular respiration, in the sensory cortex.

We extended this integral score and pathway analysis to all clusters of all regions (**Fig. 3h**). While most clusters, regardless of region, had similar integral scores, a few clusters had remarkably higher or lower delta integral scores (**Fig. 3h, delta integral annotation on heatmap**), suggesting a more prominent vulnerability or resilience, respectively, to aging. For example, Cluster 1 of the sensory cortex, representing cellular respiration, and Cluster 4 of the visual cortex, representing protein localization and synaptic signaling, had among the lowest delta integral scores (0.92 and 1.43, respectively), implicating these processes in these regions are more resilient to aging compared to the same processes or other processes of other regions. Conversely, Cluster 6 of the hypothalamus, representing protein localization and folding, and Cluster 3 of the hippocampus, representing cytoskeletal processes, had among the highest delta integral scores, suggesting these processes in these regions are more vulnerable to aging compared to the same processes or other processes of other regions. Overall, while many pathways among regions exhibit similar magnitudes of degradation alterations with age, some pathways in some regions show notable resilience or vulnerability to age, just as was observed on a protein level (**Fig. 2h**).

### Aged Neuronal Aggregome Displays Links to Synaptic Dysregulation and Proteinopathies

There are many potential causes for the age-related reduction in protein degradation that we observed, including but not limited to a reduction in lysosomal and proteasomal degradative capacities, deficits in protein transport to degradative organelles, and the formation of protein degradation-resistant protein aggregates^4^. Given the evident increase in neuronal protein aggregate number and area with age in mice and the presence of aggregates in the aged human brain as detected by Proteostat, a fluorescent dye which intercalates into beta-sheet structures of aggregated proteins found in aggresomes^31^ (**Fig. 4a, 4b, 4c**) along with the relevance of protein aggregates in age-related brain diseases^19^, we followed up on the contribution of aggregate formation to age-reduced protein degradation. Combining protein aggregate isolation techniques^32^ with neuronal BONCAT labeling enabled us to define the neuronal aggregome, the first at-scale catalog of neuronal proteins contributing to aged brain protein aggregates (**Fig. 4d**). We collected the brains of neuron-labeled aged mice, isolated the insoluble protein/aggregate fraction, and performed bead-based pull-down of azido-modified proteins to obtain the aged neuronal aggregome (**Fig. 4d**). By LC-MS, we identified 574 neuronal proteins present in aged aggregates (**Fig. 4e**). Some proteins identified are well known to aggregate in neurodegenerative diseases, including TDP-43, FUS, TMEM106b, NSF, and APP (**Fig. 4f**), and aggregation of these proteins could further perpetuate global proteostasis aberrations. Most aggregating neuronal proteins identified in the aged brain have not yet been reported or functionally examined, including the top five enriched aggregating proteins HAPLN4, HNRNPA3, PSMA4, SEC22B, PSMA2 (**Fig. 4e**). Aggregation/loss of function of these top five proteins alone, representing proteasome components (PSMA2, PSMA4), synaptic proteins (HAPLN4, SEC22B, PSMA4), or RNA-binding (HNRPNPA3), can explain both proteostasis decline and synapse dysfunction with age. Further supporting the likely relevance of these aggregating proteins in contributing to aging and diseases, 46.69% of the aggregating neuronal proteins were risk genes as defined by the H-MAGMA study (**Fig. 4f, 4g, 4h**). Several protein features implicated in aggregation propensity were altered between aggregating and non-aggregating proteins of the sensory cortex, including an increase in protein length, intrinsic unfolding propensity length (IUPL), and content of negatively charged amino acids (**Extended Data Fig. 5a**). By Gene Ontology Cellular Component analysis, aggregating neuronal proteins could be ascribed to several neuronal compartments, but the synaptic compartment was identified with the most statistical confidence and consisted of the highest number of proteins represented (**Fig. 4i**). Synapse-related terms, including cell junction, post-synapse, pre-synapse, and cell projection, recurrently appeared in the top 15 enriched cellular components (**Fig. 4i**). Synaptic proteins identified represented an array of synaptic anatomy and function (**Extended Data Fig. 5b**). Reflecting these data, Gene Ontology Biological Function analysis showed several cellular functions to be enriched, with both synaptic signaling and protein localization being recurrently represented (**Fig. 4j**).

**Figure 4:**
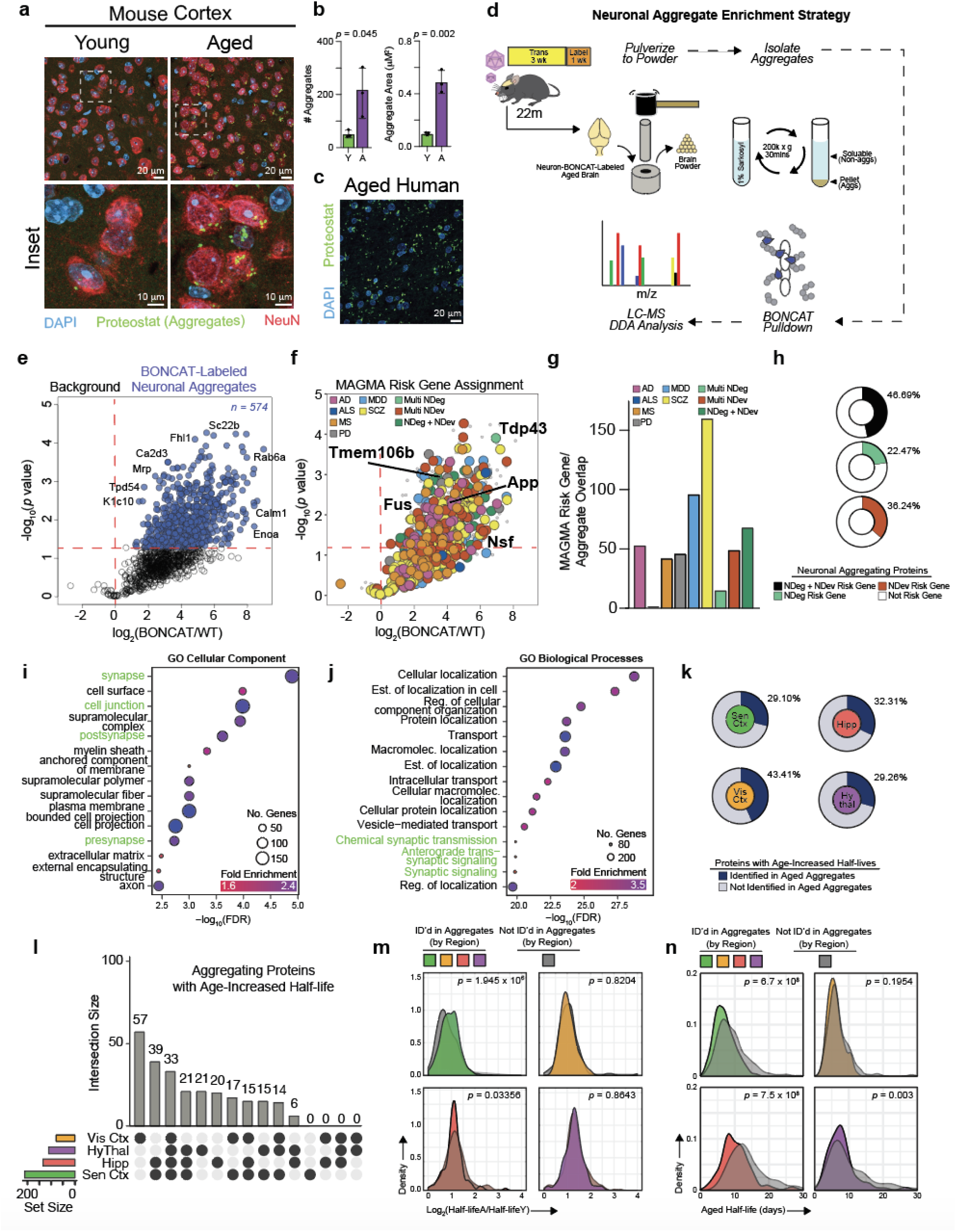
Aggregating Neuronal Proteins in Aged Brains Have Links to Age-Related Degradation Deficits, Synaptic Dysregulation, and Proteinopathies. a. Fluorescence images of young (4 months) and aged (24 months) mouse cortex tissue sections stained for neurons (NeuN, red) and protein aggregates (Proteostat, green). b. Quantification comparing aggregate number (left) and area (right) between young and aged mouse cortices. *P* value determined by two tailed t test. c. Fluorescence images of old human brain tissue section stained for protein aggregates (Proteostat, green). d. Schematic of the experimental approach to determine the identity of neuronal proteins in aggregates from the aged (22 months) mouse brain. n = 3 mice per experimental group. e. Volcano plot showing the enrichment of BONCAT-labeled neuronal proteins in aged protein aggregates identified by LC-MS relative to the wildtype background control. Proteins with a log_2_ fold change > 0 over wildtype background controls with a *p* value < 0.05 are considered hits. f. Color-coded volcano plot showing the enrichment of BONCAT-labeled neuronal proteins in aged protein aggregates identified by LC-MS relative to the wildtype background control. Proteins are color-coded based on the disease or disorder for which they have been identified as risk genes. Risk genes were derived from the H-MAGMA study and considered only if the reported *p* value was < 0.05/ g. Bar plot of the number of aggregating neuronal proteins in aged brains that are also H-MAGMA risk genes of the indicated brain diseases and disorders. Risk genes were derived from the H-MAGMA study and considered only if the reported *p* value was < 0.05. h. Donut plots showing the percentage of all aggregating neuronal proteins in aged brains that are risk genes of both neurodegenerative diseases and neurodevelopmental disorders (top), only neurodegenerative diseases (middle), or only neurodevelopmental disorders (bottom). Risk genes were derived from the H-MAGMA study and considered only if the reported *p* value was < 0.05. i. Gene Ontology Cellular Component analysis on all aggregating neuronal proteins in aged brains. Cellular component terms in green font highlight synaptic terms. Proteins used in the analysis had a log_2_ fold change > 0 over the respective background control with a *p* value < 0.05. j. Gene Ontology Biological Processes analysis on all aggregating neuronal proteins in aged brains. Cellular component terms in green font highlight synaptic terms. Proteins used in the analysis had a log_2_ fold change > 0 over the respective background control with a *p* value < 0.05. k. Donut plots showing the percentage of all proteins with an age-increased half-life in the sensory cortex (top left), visual cortex (bottom left), hippocampus (top right), and hypothalamus (bottom right) that were also identified in aged protein aggregates. l. Upset plot showing the overlap of aggregating neuronal proteins with age-increased half-lives between the indicated brain regions. m. Density plot comparing the log_2_ fold change in half-life from young to aged of aggregating neuronal proteins compared to proteins not identified as aggregated for the sensory cortex (top left), hippocampus (bottom left), visual cortex (top right), and hypothalamus (bottom right). *P* value determined by Kolmogorov-Smirnov Test. n. Density plots comparing protein half-life of aged proteins identified in aggregates compared to proteins not identified as aggregated for the sensory cortex (top left), hippocampus (bottom left), visual cortex (top right), and hypothalamus (bottom right). *P* value determined by Kolmogorov-Smirnov Test.

Cumulatively, the Gene Ontology analyses indicate that synaptic proteins are particularly vulnerable to aggregation, which coincides with evidence of age-related synaptic dysfunction and loss^27^. On a technical note, most proteins (555, or 96.69%) identified were present in two other label-free aged brain aggregate datasets (**Extended Data Fig. 5c**) and no change in the total mass of insoluble proteins (**Extended Data Fig. 5d**) or Proteostat signal (**Extended Data Fig. 5e**) was observed between BONCAT-labeled brains and non-BONCAT labeled brains, indicating BONCAT labeling does not artificially induce aggregation yet gives us the novel information about the cellular origin of aggregating proteins.

### Relationship of Aggregated Neuronal Proteins to Age-Reduced Degradation

We observed aggregation of 574 neuronal proteins in the aged brain, but questioned whether these aggregating proteins could explain the slower degradation that accompanies aging. 29.1% to 43.41% of proteins with age-reduced degradation were also found in neuronal aggregates (**Fig. 4k**). 33 proteins, including 17 synaptic proteins (VCP, HSPA8, KIF5B, PLCB1, EEF2, among others) displayed both reduced degradation with age and aggregation with age among all brain regions examined (**Fig. 4l**) and a range of proteins (6-39) displayed both reduced degradation with age and aggregation with age among 2-3 regions (**Fig. 4l**), cumulatively indicating many proteins are prone to both reduced degradation and aggregation in a non-region dependent manner. Still, each region except for the sensory cortex possessed a unique, non-overlapping set of proteins that displayed both reduced degradation with age and aggregation with age (**Fig. 4l**), potentially contributing to region-dependent vulnerability to certain aging phenotypes. While the half-life fold change of aggregating proteins in any given region was similar to that of non-aggregating proteins in the same respective region (**Fig. 4m**), the distribution of aged protein half-lives of aggregating proteins in the sensory cortex, hippocampus, and hypothalamus was slightly less than that of non-aggregating proteins in the same respective region (**Fig. 4n**), consistent with the observation that shorter-lived proteins are more aggregate prone^33^. Collectively, these data suggest that protein aggregation could be a contributor to the reduced protein degradation observed with age.

### Microglia Accumulate Neuronal Proteins with Age-Related Aberrations in Proteostasis

Microglia play critical roles in maintaining neuronal homeostasis by detecting, engulfing, and processing neuron-derived proteins^34,35^. We next aimed to identify neuron-derived proteins that accumulate in microglia seeking to infer potential neuronal-microglia interactions that contribute to neuronal proteostasis. We labeled the neuronal proteome for one week after which we freshly isolated viable CD11b+ microglia from the whole brain by fluorescence-activated cell sorting (FACS) (**Fig. 5a**), using an engulfment inhibitor cocktail^36^ throughout the process to prevent artificial *ex vivo* engulfment of neuronal proteins. We lysed the microglia and performed bead-based pull-down of any tagged neuronal proteins within them and ran the digested peptides on LC-MS (**Fig. 5a**). We identified 274 neuron-derived proteins in microglia (**Fig. 5b**). Only 18 of these proteins were predicted to contain a signal peptide (**Fig. 5c, 5d**), suggesting mechanisms of protein transfer from neurons to brain macrophages largely based on mechanisms other than the classic secretory pathway. Most proteins in our dataset were identified as mammalian exosome cargo by ExoCarta^37^ (**Fig. 5e, 5f**), but this does not exclude the possibility of proteins being transferred by other mechanisms. Most neuronal compartments, including the cell body, axons, and dendrites, were represented by the proteins identified, but synaptic and vesicle proteins were starkly enriched (**Fig. 5g**). Over 100 hundred proteins identified (36%) were annotated as synaptic (**Fig. 5g**). The enrichment of vesicle proteins could relate to the aforementioned transfer of proteins by exosomes and/or it could represent the uptake of synaptic vesicles as a byproduct of taking up synapses or parts of synapses through mechanisms such as phagocytosis or trogocytosis. SynGo analysis^38^ revealed that proteins attributed to both pre-synaptic and post-synaptic compartments were represented in our data (**Fig. 5h**), including pre-synaptic proteins ATP2B1, NAPA, NAPB, and SYT1 and post-synaptic proteins CALM1/2/3, CAMK2A/B, and GNA01. Similarly, several synaptic functions were represented, including those related to the synaptic vesicle cycle, synaptic structure modification, neurotransmitter receptor transport, and trans-synaptic signaling (**Fig. 5h**). Supporting the idea of synaptic protein accumulation by microglia, upon mining an existing dataset^39^ of freshly isolated mouse and human microglia for synaptic proteins, we found nearly 1000 mouse and 600 human proteins were annotated as synaptic by the Gene Ontology Synaptic gene set (**Fig. 5i, 5j**). The proteins represented an array of synaptic anatomy and function (**Extended Data Fig. 6a, 6b**). Synaptic proteins were nearly as abundant in average copy number in mice as non-synaptic proteins (**Extended Data Fig. 6c**), but were relatively less abundant in average copy number compared to non-synaptic proteins in humans (**Extended Data Fig. 6d**). Overlapping the 274 neuron-derived proteins identified in microglia with the total mouse and human microglia proteomes resulted in 80 and 96 proteins overlapping between our dataset and those of the mouse and human microglia proteomes, respectively (**Fig. 5j, 5k**) Thus, nearly 25% of the proteins we identified as transferred from neurons to microglia were validated in independent-label free datasets, one of which was human. We hypothesize the remaining 75% were not identified in the independent data sets because the proportion of neuron-derived proteins within the microglia proteome is relatively rare and cannot be detected by LC-MS; however, this speaks to the utility of BONCAT methodology to detect these relatively rare inter-cellularly transferred proteins.

**Figure 5:**
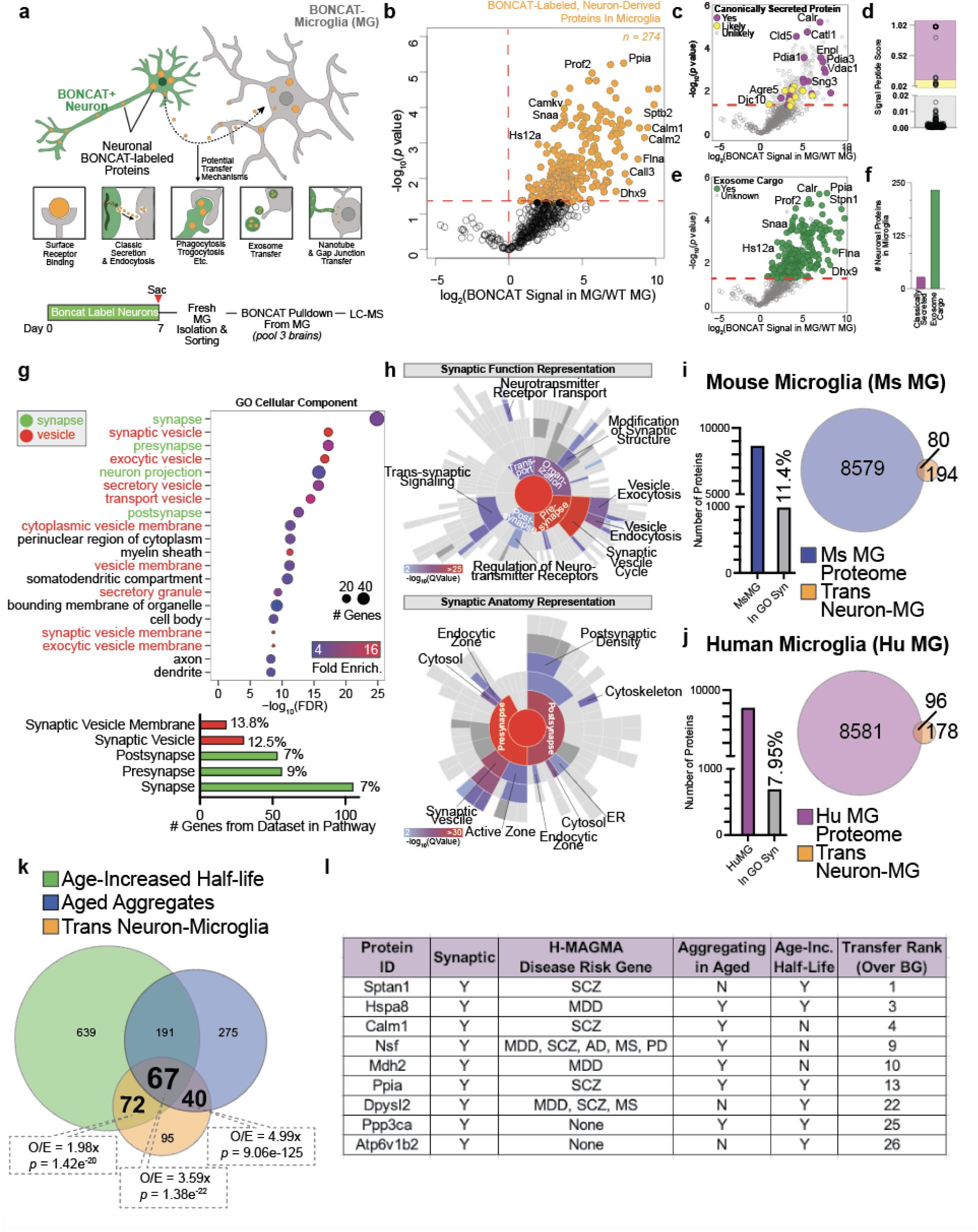
Neuronal Proteins Transferred to Microglia are Predominated by Synaptic Identity and Have Age-Related Proteostasis Aberrations. a. Schematic of the experimental approach to identify BONCAT-labeled neuronal proteins in or on microglia (MG) from young (3 months) Camk2aCre;PheRS* mice. n = 3 replicates per group with each replicate being the brains pooled from 3 mice. b. Volcano plot showing the enrichment of BONCAT-labeled neuronal proteins in microglia identified by LC-MS relative to the wildtype background control. Proteins with a log_2_ fold change > 0 over wildtype background controls with a *p* value < 0.05 are considered hits. c. Color-coded volcano plot showing the enrichment of BONCAT-labeled neuronal proteins in microglia identified by LC-MS relative to the wildtype background control. Proteins are color-coded based on the likelihood of being a canonically secreted protein as determined by signal peptide score, a score that indicates the likelihood of having a signal peptide sequence. Signal peptide scores > 0.1 are considered secreted. Signal peptide scores < 0.1 but > 0.02 are considered likely secreted. Signal peptide scores < 0.02 are considered unlikely to be secreted. d. Plot showing the signal peptide score of all neuronal proteins identified as being transferred to microglia. e. Color-coded volcano plot showing the enrichment of BONCAT-labeled neuronal proteins in microglia identified by LC-MS relative to the wildtype background control. Proteins are color-coded based on being reported as exosome cargo by Exocarta. f. Bar plot of the number of neuronal proteins transferred to microglia that are either classically secreted or reported exosome cargo. g. Gene Ontology Cellular Component analysis on all neuronal proteins transferred to microglia. Proteins used in the analysis had a log_2_ fold change > 0 over the respective background control with a *p* value < 0.05 (top). Bar chart of the number of proteins identified in the dataset that contribute to the indicated gene ontology terms and the percent of the gene list represented by the identified proteins (bottom). h. Sunburst plots showing synaptic functional representation (top) and synaptic anatomical representation (bottom) of neuronal proteins identified in microglia. i. Bar plot of the number of proteins identified in the mouse microglia proteome and the number of those proteins within the Gene Ontology Synapse gene list (left) and Venn Diagram showing the overlap of proteins identified in the entire mouse microglia proteome and the neuron-derived proteins we identified in microglia. j. Bar plot of the number of proteins identified is the human microglia proteome and the number of those proteins within the Gene Ontology Synapse gene list (left) and Venn Diagram showing the overlap of proteins identified in the entire human microglia proteome and the neuron-derived proteins we identified in microglia. k. Venn Diagram showing the overlap of neuronal proteins identified to have a reduction in degradation with age (green), identified in aged aggregates (blue), and transferred to microglia (yellow). Over enrichment of overlapping proteins and associated p values derived from hypergeometric test. Values for two-way overlap comparisons based on total overlap between the datasets, not only the number indicated in the Venn Diagram, which does not include the 67 proteins overlapping between all three datasets. l. Table of information on selected proteins transferred from neurons to microglia and also present in aged aggregates and/or display an increased half-life with age. Y = yes, n = no.

Lastly, we questioned whether any of the neuronal proteins that accumulated in microglia were those observed to have age-related proteostasis aberrations. Overlapping the lists of proteins identified to have slower degradation kinetics with aging (**Fig. 2**), proteins present in aged neuronal aggregates (**Fig. 4**), and neuronal proteins found in microglia (**Fig. 5**) resulted in 67 overlapping proteins (**Fig 5k, 5l**). 72 proteins overlapped uniquely between slower degradation kinetics with aging and neuronal proteins found in microglia and 40 proteins overlapped uniquely between aged neuronal aggregates and neuronal proteins found in microglia (**Fig. 5k, 5l**). Cumulatively, 179 of the 274 (65.33%) neuronal proteins found in microglia showed age-related proteostasis deficits, whether age-reduced degradation or presence in aggregates. Each intersection of protein lists had an over-enrichment of overlapping proteins beyond which would be expected by random chance (1.98x for age-increase half-life/neuron-microglia transfer intersection; 4.99x for aged aggregate/neuron-microglia transfer intersection; 3.59x for three-way intersection) (**Fig. 5k**). In addition to many proteins being synaptic, several of these proteins, such as NSF, DPYSL2, SPTAN1, and MDH2 are brain disease risk genes (**Fig. 5l**).

From these results, we conclude the accumulation of neuronal proteins with age-related alterations (slower degradation and/or aggregation) in microglia is not random but rather a specific mechanism to remove aberrant protein species from neurons to maintain neuronal proteostasis (**Extended Data Fig. 6e**).

## Discussion

Loss of proteostasis is a hallmark of aging with crucial implications for neurons and thus cognitive function and dementia risk^1,2,4^. Despite efforts to understand brain proteostasis with aging^23,24^, no studies have been performed at scale *in vivo* with a neuron-specific perspective. Here, aided by new nascent proteome labeling models that enable cell-specific at-scale analysis of proteostasis dynamics in mice across lifespan, we were able to study key aspects of neuronal proteostasis with aging, namely, protein degradation, protein aggregation, and inter-cellular protein transfer of neuronal proteins to microglia. We first showed that neuronal protein degradation rates decline on average by approximately two-fold with aging, a deficit emerging mostly after middle age and showing region-dependent variations in relation to the extent of turnover reduction. The reduction in neuronal protein degradation is consistent with two studies examining age-related protein turnover changes at the whole brain and synaptosome levels using SILAC pulse-chase methodology^23,24^. However, we observed on average a doubling in protein half-life with age, while a study examining whole-brain protein degradation reported a 20% average increase^24^, a difference at least partially explained by the neuron-specific perspective achieved in our study or the difference in timepoint sampling. By examining neuronal proteins that aggregate in aged brains, we found approximately 30% of neuronal proteins with age-reduced degradation also aggregated with age, suggesting aggregation is one likely contributor to reduced degradation, albeit not the singular cause, and it cannot be excluded as a consequence of reduced degradation. A common theme emerged among proteins displaying age-reduced degradation and the propensity to aggregate in that there was an enrichment for synaptic proteins representing a diverse array of synaptic compartments and functions. This is intriguing considering evidence of synaptic dysfunction and loss with aging and age-related diseases, suggesting that loss of synaptic proteostasis could be at the center of age-related synapse dysfunction. Given many of these proteins are neurodegenerative risk genes, it is likely some or all risk gene variants of the proteins we identified confer a propensity for altered degradation or aggregation, just as has been observed of mutations in proteins including APP, TDP-43, HTT, and SOD1.

Proteostasis is traditionally described as being intrinsically regulated within cells via chaperones, the ubiquitin-proteasome system, and autophagy^4^. While these mechanisms are key, emerging evidence suggests that cell-extrinsic partners also play a role. For example, transsynaptic transfer of protein aggregates between neurons can help balance proteostasis across cells^40^. We hypothesized that neuronal proteostasis is partially maintained by the transfer of neuronal proteins to microglia, cells that are known to contribute on a broader level to brain homeostasis^34,35,41^. Enabled by our protein labeling models, we identified 274 BONCAT-labeled neuronal proteins that accumulated in microglia, which were enriched for synaptic proteins.

Intriguingly, 65% of these proteins also displayed age-related proteostasis aberrations, having slower degradation with age and/or being found in aged neuronal aggregates. The number of age-impaired proteins that accumulate in microglia is significantly more (minimally two-times more) than expected by random chance, thus we propose that the transfer of these protein species from neurons to microglia is a mechanism to maintain neuronal proteostasis. Indeed, there is mounting evidence for intercellular spread of protein aggregates from neurons to glia via their release through exosomes or transfer through tunneling nanotubules^42–44^. As synaptic proteins are enriched among the neuronal proteins transferred to microglia as well as the proteins found in brain aggregates, we hypothesize an additional mechanism may be the selective engulfment of proteostatically-stressed synapses by microglia. This mechanism may partially explain synaptic dysfunction and loss with age^27,45,46^. Besides better understanding the mechanisms of protein transfer, the effects of the transfer on neurons, microglia, and the brain as a whole will be imperative to explore. The transfer of old and aggregated proteins from neurons to microglia may represent a short-term gain to neurons but a long-term loss when considering the combined detrimental effects these proteins could have on recipient microglia and the collective loss of synapses. Cumulatively, our findings reveal several age-related neuronal proteostasis aberrations that have links to synaptic dysregulation and proteinopathies, putting forward new hypotheses related to the causes of age-related synaptic dysfunction. It will be imperative for future studies to elucidate the consequences of declined neuronal proteostasis and develop therapies to restore neuronal proteostasis to promote resilience to brain aging and diseases.

## Acknowledgements

We thank all members of the Wyss-Coray Lab, with special thanks to past members Kyle Brewer and Steven Shuken, for insightful comments provided through the genesis of this work and Divya Channappa, Kathleen Dickey, and Hui Zhang for laboratory management. We also thank members of the Carolynn Bertozzi Lab, namely Nicholas Riley, Brendan Floyd, and Johannes Florian Hevler for consultation regarding LC-MS experimental design and analyses. We gratefully acknowledge the contribution of Kim Martens and the Genetic Engineering Technologies Service at the Jackson Laboratory. NIH Pathway to Independence Award 1K99AG088304-01 (I.H.G), MAC3 Dementia and Ageing Fellowship (I.H.G., N.L.), DFG Compute and Storage Cluster 469073465 (A.K.), US Department of Defense HT9425-23-1-0879 (J.L.), NIH AG059694 (J.L.), NIH Director’s Early Independence Award 1DP5OD033381 (A.C.Y), Burroughs Welcome Fund Career Awards at the Scientific Interface (A.C.Y.), Stanford Bio-X Fellowship (S.M.S.), AHA-Allen Brain Health and Cognitive Impairment Cross-Network Collaborative Grants 23BHCICG1188316 (N.L.). Novel knock-in mouse models were developed with funding from an anonymous organization.

## Author Contributions

I.H.G. and T.W-C. conceptualized the project. I.H.G. performed most of the experiments with the assistance described as follows: P.S.W and N.T. assisted in running samples on LC-MS under the supervision of T.C.; K.C., N.L., Z.G., and J.L. assisted in some mouse procedures and/or collections; K.C. and Z.G. assisted with in-gel fluorescence and/or tissue staining experiments and steps of azido-modified protein bead-based pull downs. A.C.Y., M.S., and T.W-C. created PheRS* and TyrRS* knock-in mice. D.G. performed molecular analysis of PheRS* and TyrRS* knock-in founders and developed genotyping assays. I.H.G. created BONCAT AAV expression constructs. I.H.G. devised azido-modified amino acid administration protocols. I.H.G., P.S.W., and N.T. processed LC-MS data. I.H.G., V.P.W., P.M-L., and B.T.M. performed formal analyses, with analyses related to protein degradation kinetics, half-life, and clustering being under the supervision of A.K.. I.H.G., V.P.W, P.M-L., B.T.M., H.O., Y.G., A.K., A.C.Y., and T.W-C. developed data analysis methodology. I.H.G., S.M.S., and A.C.Y. developed wet bench methodology related to click chemistry. I.H.G. and T.W-C. wrote the original draft. T.W-C. supervised the study.

## Additional Information

Correspondence and requests for materials should be addressed to Tony Wyss-Coray.

Reprints and permissions information is available at www.nature.com/reprints

## Methods

### Mouse Husbandry

Mice were housed in standard conditions on a 12-hour light-dark cycle and provided water and standard chow *ad libitum*. In some experiments, as documented in the main text, mice were provided with azido-amino acid infused water or 0.1% methionine, 0.35% cysteine (low methionine) chow (Envigo, TD.160659). All animal procedures were approved by the Administrative Panel on Laboratory Animal Care at Stanford University. All experiments used male mice.

### Mouse Sources

All transgenic mouse lines were obtained from The Jackson Laboratory (Bar Harbor, Maine). Besides the BONCAT lines generated in this study, the generation of which will be described below, the transgenic lines used in this study include B6.Cg-Tg(Camk2a-cre)T29-1Stl/J (The Jackson Laboratory, 005359); B6.C-Tg(CMV-cre)1Cgn/J (The Jackson Laboratory, 006054); C57BL/6-Gt(ROSA)26Sor^tm1(CAG-GFP,-Mars*L274G)Esm/J^ (The Jackson Laboratory, 028071); B6;C3-Tg(Prnp-MAPT*P301S)PS19Vle/J (The Jackson Laboratory, 008169). Homozygous cre lines were bred to homozygous BONCAT lines to generate offspring heterozygous for the cre driver and heterozygous for the BONCAT transgene. The B6;C3-Tg(Prnp-MAPT*P301S)PS19Vle/J line was maintained by crossing hemizygous males to non-carrier females, with hemizygous mice being used as experimental mice and aged-matched non-carriers being used as wildtype controls. Wildtype C57BL/6 mice used for aging-related AAV transduction experiments were obtained from the National Institute of Aging colony. Wildtype C57BL/6 used for non-aging related AAV transduction experiments or used as background controls were obtained from The Jackson Laboratory.

### Transgenic Mouse Generation

The new BONCAT models, PheRS^T413G^ and TyrRS^Y43G^, introduced in this manuscript were generated in collaboration with The Jackson Lab. sgRNAs (ACTGGAGTTGCAGATCACGA and GCAGATCACGAGGGAAGAGG) were designed to insert a cassette encoding a CMV-IE enhancer/chicken beta-actin/rabbit beta-globin hybrid promoter (CAG) followed by a floxed STOP cassette containing 3xSV40 polyadenylation signals, an EGFP sequence, a viral 2A oligopeptide (P2A) self-cleaving peptide, that mediates ribosomal skipping, and either the PheRS^T413G^ gene or the TyrRS^Y43G^ gene into the Gt(ROSA)26Sor locus. gRNA, the cas9 mRNA, and a donor plasmid were introduced into the cytoplasm of C57BL/6J-derived fertilized eggs with well recognized pronuclei. Injected embryos were transferred to pseudopregnant females.

Surviving embryos were transferred to pseudopregnant females. Resulting progeny were screened by DNA sequencing to identify correctly targeted pups, which were then bred to C57BL/6J mice for germ line transmission. This colony was backcrossed to C57BL/6J mice for at least 3 generations. Sperm was cryopreserved at The Jackson Lab. To establish our live colony, an aliquot of frozen sperm was used to fertilize C57BL/6J oocytes. Upon publication, the PheRS^T413G^ model (C57BL/6J-*Gt(ROSA)26Sor^em2(CAG-GFP,-Farsa*T413G)Msasn^*/J) and the TyrRS^Y43G^ is) C57BL/6J-*Gt(ROSA)26Sor^em3(CAG-GFP,-Yars1*Y43G)Msasn^*/J) will be made available for purchase from The Jackson Laboratory.

### Biological Replicates

All mice were maintained as individual biological replicates except for the protein aggregation experiments and neuron-to-brain macrophage protein transfer experiments. In the two aforementioned experiments, isolated labeled neuronal protein from aggregates or brain macrophages was expected to be relatively minimal, so to ensure detection by LC-MS, the entire brain of three mice was pooled to generate one biological replicate.

### Mouse AAV Injection

Mice were anesthetized with Isoflurane and 3e11-5e11 AAV genome copies were injected in 100 μL sterile 1x PBS via the retroorbital sinus. Equal genome copies were injected in mice between which comparisons would be made. For maximal transgene expression, mice were used no sooner than 3 weeks following initial transduction.

### Non-canonical Amino Acid Preparation and Administration

All azido-modified amino acids (AzAAs), including 4-Azido-L-phenylalanine (Vector Laboratories; 1406-5G), N-epsilon-Azido-L-lysine hydrochloride (Iris Biotech, HAA1625.0005), and 3-Azido-L-tyrosine (Watanabe Chemical Industries, J00560) were prepared as a 12.35 mg/mL solution for intra-peritoneal injections. Unless otherwise noted in this manuscript, AzF was injected once daily for 7 consecutive days.

For the experiment providing N-epsilon-Azido-L-lysine hydrochloride-infused water to mice, N-epsilon-Azido-L-lysine hydrochloride was dissolved in sterile mouse water to a final concentration of 30 mM, a concentration reported to provide excellent Camk2a+ neuronal labeling. The water was brought back to its original pH by addition of NaOH. For the experiments providing 4-Azido-L-phenylalanine-infused water to mice, 4-Azido-L-phenylalanine was dissolved in sterile mouse water at 1 mg/mL. The water was brought back to its original pH by addition of NaOH.

### Tissue Harvesting and Handling

Mice were anesthetized with Isoflurane and transcardially perfused with at least 20 mL of 1x PBS. After perfusion, brains were immediately extracted and either fixed in 4% PFA, snap-frozen in tubes on dry ice, or immediately enzymatically dissociated. For experiments requiring brain region dissection, after brain extraction, regions were immediately dissected on ice using a ‘rodent brain matrix’ 1-mm coronal slicer (Tedpella, 15067) according to coordinates obtained from the Mouse Brain Library (http://www.mbl.org/) using the C57BL/6J atlas as reference. Upon dissection, regions were snap-frozen in tubes. In cases in which tissue was fixed, tissue was fixed for 24 hours and then prepared for either paraffin embedding or sucrose cryoprotection. Snap-frozen tissue was stored long-term at −80C.

### Microglia Isolation

To isolate microglia, whole brains were first enzymatically dissociated as previously described^1^ with the addition of an engulfment inhibitor cocktail to prevent *ex vivo* engulfment of neuronal debris^2^. At all steps during microglia isolation, staining, and sorting, liquids were supplemented with the engulfment inhibitor cocktail containing the final concentrations of the following reagents: 25 µM Pitstop2 (Abcam, ab120687), 2 µM Cytochalasin D (Tocris, 12330), 2 µM Wortmannin (Tocris, 1232), 40 µM Dynasore (Tocris, 2897), 40 µM Bafilomycin A1 (Tocris, 1334), with each reagent being prepared as a 1000x stock. Immediately after extracting perfused brains, they were placed in 800 µL 1x D-PBS+/+ (Thermo Fisher Scientific, 14040117) on ice. Next, brains were minced on ice using fine scissors for approximately two minutes until brain chunks were small enough to triturate with a p1000 pipette with little resistance during pipetting. Brains were triturated until there was no resistance during pipetting. Brain suspensions were pelleted by centrifugation at 300g for 5 minutes at 4C. The supernatant was removed by pipetting and an enzymatic cocktail prepared from Multi-tissue Dissociation Kit 1 (Miltenyi Biotec, 130-110-201), consisting of 100 µL Enyzme D, 50 µL Enyzme R, 12.5 µL Enyzme A, and 2.4mL D-PBS+/+, was added. The pellet was resuspended by pipetting, after which the suspensions were transferred to a tube rotator at 37C for a 20 minute incubation. Halfway through and at the end of this incubation, brain suspensions were triturated with a p1000 approximately 20 times to help break up brain chunks until the suspension was largely void of any visible chunks. After the incubation, 10 mL ice cold DPBS+/+ was added to each brain suspension, after which the entire suspension was run through a 70 µm cell strainer into a new tube. Filtered suspensions were centrifuged at 500g for 10 minutes at 4C and the supernatant removed by pipetting. Next, myelin was removed from the preparations using Debris Removal Solution (Miltenyi Biotec, 130-109-398). Each brain pellet was resuspended up to 3.1 mL with cold D-PBS+/+ and 0.9 mL Debris Removal Solution was added to each pellet and mixed by gentle inversion of the tube. 4 mL cold D-PBS+/+ was overlaid on top of the brain/Debris Removal Solution mixture. Samples were centrifuged at 3000g for 13 minutes at 4C with medium acceleration and 0 break. Following centrifugation, the myelin interface and liquid above it were removed by pipetting and 11 mL of cold DPBS+/+ was mixed with the remaining cell suspension. The cell suspensions were centrifuged at 1000g for 13 minutes at 4C with 0 break, and the resulting supernatant removed by pipetting. The largely myelin-depleted cell pellets were resuspended in 80 µL AstroMACS Separation Buffer (Miltenyi Biotec, 130-117-336) containing 10 µL FcR Blocking Reagent, mouse (Miltenyi Biotec, 130-092-575) and incubated on ice for 10 minutes. Next, 10 µL anti-ACSA-2 MicroBeads (Miltenyi, 130-097-679) was mixed in to the cell suspension and incubated for 15 minutes on ice. After incubation, cells were was in 1 mL AstroMACS Separation Buffer and centrifuged at 300g for 10 minutes at 4C. The supernatant was removed and pellets were resuspended in 500 µL AstroMACS Separation Buffer and loaded onto a pre-washed LS Column (Miltenyi Biotec, 130-042-401). The LS Columns were washed three times, each time with 3 mL AstroMACS Separation Buffer. The flow through was retained, as this contained microglia, while the cells retained in the column were eluted with 5 mL of AstroMACS Separation Buffer and retained as an astrocyte fraction used for other experiments. The cell suspensions were washed by centrifugation at 300g for 10 minutes at 4C. Supernant was removed and cells were stained 1:10 with APC/Cyanine7 anti-mouse/human CD11b Antibody (BioLegend, 101225) and Calcein-AM (BioLegend, 425201) at a final concentration of 1x in Cell Staining Buffer (BioLegend, 420201) for 30 minutes on ice. Cells were washed in 1 mL Cell Staining Buffer by centrifugation at 300g for 10 minutes at 4C. Supernatant was removed and cells were resuspended in an appropriate volume of Cell Staining Buffer for Fluorescence Activated Cell Sorting (FACS). CD11b+/brain macrophages were sorted by gating on CD11b+, Calcein-AM+ singlets on a Sony MA900 cell sorter (Sony Biotechnology, Inc). Three biological replicates-worth of brain macrophages were pooled into a single replicate and frozen at −80C before lysing the cells and enriching for BONCAT-labeled proteins.

### Cloning of *Mm*PheRS_T413G_ into AAV Vector

The sequences for PheRST_413G_ and Camk2a promoter were ordered as gBlocks HiFi Gene Fragment from IDT (Integrated DNA Technologies, USA). The sequences were derived from prior publications^3,4^ and modified only to add a 3x Flag Tag to the c-terminus of PheRST_413G_ as well as flanks to both fragments to enable Gibson Assembly cloning.

The PheRST_413G_ gene fragment was first cloned into pAAV-CAG-GFP (Addgene, 37825) backbone by removal of GFP by BamHI/EcoRV double restriction digest followed by Gibson Assembly (NEB, E2611S). From this Gibson Assembly product, we then removed the CAG promoter by XbaI/NdeI double restriction digest and cloned in the Camk2a promoter by Gibson Assembly. Propagation of AAV plasmids was performed in NEB Stable Competent E. coli (NEB, C3040H) at 30C to avoid mutations in the AAV ITR sequences.

### AAV Production

AAV was custom-produced by VectorBuilder (VectorBuilder, Inc, Chicago, IL). All preparations were ultra-purified and utilized PHP.eB serotype.

### Copper-Catalyzed Click Reaction on Lysates for In-Gel Fluorescence

Brain tissue was first homogenized by sonication in a strong lysis buffer (8 M Urea, 1%SDS, 100 mM choloracetamide/CAA, 20 mM Iodoacetamide/IAA, 1 M NaCl, and 1x protease inhibitor in 1x PBS). Sonication was performed for at least three cycles of 10 seconds sonicating with at least 5-second breaks between sonication cycles at an amplitude of 90% using a probe sonicator. Homogenates were centrifuged for 15 minutes at >16,000g at 4C. The resultant supernatant was retained, and aliquots were immediately measured by BCA to obtain protein concentration with the remaining supernatant being frozen at −80C until it was further processed to enrich for BONCAT-labeled proteins.

Samples to be compared were normalized to equal protein amounts (110 mg) and brought up to 33.3 µL total with water. A click reaction was performed on the lysates to ‘click’ a fluorophore onto azide side chains of labeled proteins. The following chemical cocktail was added to the normalized lysates for one hour with constant shaking to perform the click reaction: 0.83 µL Alexa Fluor 647 Alkyne, Triethylammonium Salt (Thermo Fisher Scientific, A10278)at 5 mM, 1.04 µL Copper (II) Sulfate (Millipore Sigma, 451657-10G) at 6.68 mM, 2.087 µL THPTA (Click Chemistry Tools, 1010-500) at 33.3 mM, 4.17 µL Aminoguanidine hydrochloride (Millipore Sigma, 396494-25G) at 100 mM, 8.33 µL Sodium L-Ascorbate (Fisher Scientific, A0539500G) at 100 mM, 33.5 µL PBS. Importantly, 20 mM CuSO4 and 50 mM THPTA were mixed at a 1:2 ratio for 15 minutes before combining the rest of the click reaction. After the one hour click reaction incubation, the reactions were filtered through Zeb Spin Desalting Columns, 7K MWCO, 0.5 mL format (Thermo Fisher Scientific, 89882) following the manufacturer’s protocol to remove unbound fluorophore. The flow through containing the clicked lysates was retained. 21 µL of the clicked lysates was mixed with 7 µL 1x loading buffer, which was prepared by mixing 10 µL 2-Mercaptoethanol (Millipore Sigma, M6250-100ML) with 115 µL 4x NuPAGE LDS Sample Buffer (Thermo Fisher Scientific, NP0007). Samples were heated at 95C for 10 minutes to denature the proteins. Clicked and denatured lysates were loaded onto a NuPAGE 12%, Bis-Tris gel (Thermo Fisher Scientific, NP0341BOX) and run at 200V for 45 minutes. The gel was imaged to detect the Alexa 647-clicked proteins using the LI-COR Odyssey XF imaging system. To detect total loaded protein, gels were stained with GelCode Blue Stain Reagent (Thermo Fisher Scientific, 24590), destained in water for at least one hour, and then again imaged using the LI-COR Odyssey XF imaging system.

### Copper-Catalyzed Click Reaction for Tissue Fluorescence Microscopy

Tissue sections were prepared for click staining of azide-modified proteins as described in the immunofluorescence staining of tissue sections subsection through the blocking step. After blocking, tissue sections were stained for one hour in the following click reaction cocktail: 2 µL Alexa Fluor 647/594/555/488 Alkyne, Triethylammonium Salt (Thermo Fisher Scientific, A10278) at 5 mM, 5 µL Copper (II) Sulfate (Millipore Sigma, 451657-10G) at 20 mM, 10 µL THPTA (Click Chemistry Tools, 1010-500) at 50 mM, 100 µL Aminoguanidine hydrochloride (Millipore Sigma, 396494-25G) at 50 mM, 100 µL Sodium L-Ascorbate (Fisher Scientific, A0539500G) at 50 mM, 783 µL PBS. Importantly, 20 mM CuSO4 and 50 mM THPTA were mixed at a 1:2 ratio for 15 minutes before combining the rest of the click reaction. After staining, tissue sections were washed three times in tris-buffered saline-Tween 20 (TBS-T) and either stained further with antibodies as described in the immunofluorescence staining of tissue sections subsection or mounted and coverslipped.

### Immunofluorescence Staining of Tissue Sections

For sucrose-cryopreserved tissues, 40 µm-thick sections were sectioned on a Lecia sliding microtome equipped with a cooling unit and cooling stage. In rare instances in which antigen retrieval was necessary, such as for anti-GFP staining, tissues were incubated in SignalStain Citrate Unmasking Solution (Cell Signaling Technology, #14746) diluted to 1x in distilled water for one hour at 95C. After cooling to room temperature, tissues were blocked and permeabilized in 5% normal donkey serum (Jackson Immuno Research, 017-000-121) and 0.3% Triton X-100 (Millipore Sigma, 93443-100ML) in 1x PBS for one hour. After blocking, tissues were stained with primary antibody diluted in 1% (W/V) bovine serum albumin (Fisher Scientific, BP9703100) and 0.3% Triton X-100 in 1x PBS overnight at 4C with gentle rocking agitation.

Primary antibodies were used at the following dilutions: rabbit anti-GFP (Cell Signaling Technology, 2956S) at 1:400; Rb anti-Iba1 (Fujifilm Wako, 019-19741) at 1:2000; guinea pig anti-NeuN (Synaptic Systems, 266 004) at 1:1000. After primary antibody staining, tissue sections were washed three times in tris-buffered saline-Tween 20 (TBS-T). After washing, tissue sections were stained with secondary antibodies diluted 1:500 in 1% (W/V) bovine serum albumin and 0.3% Triton X-100 in 1x PBS for three hours at room temperature with gentle rocking agitation. All secondary antibodies recognized the IgG domain of primaries, were conjugated to Alexa fluorophores, and purchased from Jackson Immuno Research. After secondary antibody staining, tissues were washed as described above, briefly stained with 4’,6-Diamidino-2-Phenylindole, Dihydrochloride (Thermo Fisher Scientific, D1306), and mounted and coverslipped on Fisherbrand Superfrost Plus Microscope Slides (Fisher Scientific, 12-550-15) with Fluoromount-G Slide Mounting Medium (Fisher Scientific, 50-259-73).

Staining of formalin-fixed paraffin-embedded tissue sections was similar to the methodology employed for free-floating sucrose cryopreserved tissue sections, except (1) tissue sections on slides were deparaffinized and rehydrated by incubation through a gradient of xylene, 100% ethanol, 90% ethanol, 80% ethanol, 70% ethanol, and water and (2) heat-induced antigen retrieval was always performed by heating slides in 1x SignalStain Citrate Unmasking Solution for 10 minutes in a microwave.

Proteostat aggresome staining was only performed on formalin-fixed paraffin-embedded tissue sections following deparaffinization, rehydration, antigen retrieval, and blocking of tissue sections. Proteostat (Fisher Scientific, NC0098538) was diluted 1:1000 in 1x PBS and incubated on tissue sections for 5 minutes at room temperature, and subsequently washed several times with TBS-T. After washing, slides were either coverslipped or put through antibody staining.

### Microscopy

Images of 5 µm thick FFPE tissue were captured on a Zeiss Axioimager (Zeiss). Images of mounted free-floating 40 µM thick tissue were captured on a Zeiss LSM 900 (Zeiss). When capturing images to be compared, all imaging parameters and post-acquisition processing parameters were kept identical between images to be compared.

### Fluorescence Image Analysis

Quantification of protein aggregate number and area was performed using FIJI. Images were uploaded and scale set according to image scale bar. Brightness and contrast were adjusted equally among all images. Images were converted to 8-bit and binary, after which masked particles, which represented aggregates, were analyzed and summary statistics related to aggregate number and average area were recorded.

### Western Blot

Brain tissue was first homogenized by sonication in a strong lysis buffer (8 M Urea, 1%SDS, 100 mM choloracetamide/CAA, 20 mM Iodoacetamide/IAA, 1 M NaCl, and 1x protease inhibitor in 1x PBS). Sonication was performed for at least three cycles of 10 seconds sonicating with at least 5-second breaks between sonication cycles at an amplitude of 90% using a probe sonicator. Homogenates were centrifuged for 15 minutes at >16,000g at 4C. The resultant supernatant was retained, and aliquots were immediately measured by BCA to obtain protein concentration with the remaining supernatant being frozen at −80C until it was further processed.

Samples to be compared were normalized to equal protein amounts and brought up to equal volumes with water. Samples were denatured and reduced by adding 4x NuPAGE™ LDS Sample Buffer to 1x and 2-Mercaptoethanol to 20% and heating at 95C for 5 minutes. Denatured and reduced samples were run on a NuPAGE 12%, Bis-Tris gel for approximately two hours at 110v. Proteins were transferred to 0.45 µM methanol-activated PVDF membrane by standard wet transfer methodology at 400mA for approximately 90 minutes at 4C. Membranes were blocked in Intercept TBS Blocking Buffer (Fisher Scientific, NC1660550) for one hour with gentle shaking. After blocking, membranes were incubated in primary antibody diluted 1:1000 in 5% Bovine Serum Albumin overnight at 4C with gentle shaking. Primary antibodies used are rabbit anti-beta-actin (Cell Signaling Technology, 4970S) and rabbit anti-HSP90 (Cell Signaling Technology, 4877T). Following primary antibody staining, membranes were washed three times with TBS-T with gentle shaking. Following washes, membranes were stained with IRDye 800CW Goat anti-Rabbit IgG Secondary Antibody (Li-Core, 926-32211) diluted 1:5000 in 5% Bovine Serum Albumin by light-protected incubation with gentle shaking for one hour.

Following secondary antibody staining, membranes were washed three times with TBS-T with gentle shaking. Lastly, membranes were imaged using the LI-COR Odyssey XF imaging system.

### Insoluble Protein/Aggregate Isolation

The protocol for insoluble/aggregated protein isolation was slightly modified from a previous publication^5^. Each hemisphere was ground to a powder on a liquid-nitrogen cooled pestle. The resultant homogenate powder was quickly transferred to tubes on dry ice, pooling three pulverized brains to generate one biological replicate. This methodology of cell lysis is used to preserve aggregates, which could be compromised using other lytic techniques (for example, sonication or detergent-based solutions). The homogenate powder was quickly weighed to avoid thawing. Homogenate was resuspended at 1 g/5 mL in 50 mM HEPES, 250 mM sucrose, 1 mM EDTA, and 1x Protease Inhibitor in water on wet ice. Next, for every 0.8 mL homogenate solution, 100 μL 5 M NaCl and 100 μL 10% Sarkosyl was added to the homogenate solution on wet ice. The homogenate was gently sonicated for 3 separate intervals for 5 seconds at an amplitude of 30% using a probe sonicator (Qsonica, Q125-110) at 4C. The protein concentration of the resultant bulk homogenate was determined by BCA and the bulk homogenate was frozen at −80C until further processing as described next. Equal protein amounts of bulk protein homogenate were aliquoted from each sample and diluted to 10 mg/mL in 1% Sarkosyl, 0.5 M NaCl, and 1x Protease Inhibitor in the low salt buffer used above. Samples were ultra-centrifuged at 180,000g for 30 minutes at 4C during which soluble/non-aggregated proteins remained in the supernatant and insoluble/aggregated proteins were pelleted. The supernatant was gently removed and frozen while pellets were washed in 1% Sarkosyl and ultra-centrifuged again at 180,000g for 30 minutes at 4C. Pellets were retained and frozen at −80C until further processing for various analyses. It is important to note that downstream enrichment of BONCAT-labeled proteins from total aggregates should only result in obtaining neuronal proteins that were part of aggregates, but not non-neuronal co-aggregating proteins. The reason for this is due to the buffer required for pull-down (see section below on enrichment of BONCAT-labeled proteins), which both solubilizes and denatures proteins. Introduction of total aggregates to this buffer should result in the release of non-neuronal co-aggregating proteins from neuronal aggregates. The pull-down then selectively enriches for BONCAT-labeled proteins, excluding non-BONCAT-labeled (non-neuronal coaggregating proteins) from further analysis.

### Enrichment of BONCAT-Labeled Proteins and Preparation for LC-MS

Brain tissue was first homogenized by sonication in a strong lysis buffer (8 M Urea, 1%SDS, 100 mM choloracetamide/CAA, 20 mM Iodoacetamide/IAA, 1 M NaCl, and 1x protease inhibitor in 1x PBS). Sonication was performed for at least three cycles of 10 seconds sonicating with at least 5-second breaks between sonication cycles at an amplitude of 90% using a probe sonicator. Homogenates were centrifuged for 15 minutes at >16,000g at 4C. The resultant supernatant was retained, and aliquots were immediately measured by BCA to obtain protein concentration with the remaining supernatant being frozen at −80C until it was further processed to enrich for BONCAT-labeled proteins.

Samples to be compared were normalized to equal protein amounts (1-2 mg total) and equal volumes in lysis buffer. Lipids were removed by the addition of 10 µL Cleanascite (Fisher Scientific, NC0542680) per 40 µL homogenate and incubation with constant agitation on a thermomixer set to 1500-2000rpm for 10 minutes. Samples were then centrifuged at >16,000g for 3 minutes to pellet lipids. The resultant supernatant was retained and 7.5 units of Benzonase (Millipore Sigma, 70664-3) was added per 40 µL sample to digest nucleotides over 30 minutes with constant agitation on a thermomixer set to 1500-2000rpm. Following Benzonase treatment, samples were diluted to 1mL total with lysis buffer and added to 200 µL dry control agarose beads (Thermo Fisher Scientific, 26150) pre-washed three times prior to sample addition (washed one time with water and two times with 0.8% SDS). Samples were pre-cleared with the control agarose beads to remove non-specific bead binders by one hour of light-protected end-over-end rotation. After pre-clearing, samples were centrifuged at 1,000g for 5 minutes to pellet the plain agarose beads. The resultant supernatant was added to 20 µL dry DBCO beads (Vector, 1034-25) pre-washed four times prior to sample addition (washed one time with water and three times with 0.8% SDS). BONCAT-labeled protein enrichment with the DBCO beads was performed overnight with light-protected end-over-end rotation. After overnight enrichment of BONCAT-labeled proteins to the DBCO agarose beads, 10 µL of 100 mM ANL (Iris Biotech, HAA1625.0005) was added to each sample to quench the DBCO beads to prevent further protein binding. Quenching was performed for 30 minutes with light-protected end-over-end rotation.

After quenching of the DBCO beads, samples were centrifuged for 5 minutes at 1000g and the supernatant was discarded while the DBCO beads were retained. DBCO beads were washed by the addition of 1 mL water and again centrifuged for 5 minutes at 1000g. The supernatant was discarded and 0.5 mL 1 mM dithiothreitol (Thermo Fisher Scientific, R0861) was added to each sample. Samples in 1 mM DTT were incubated for 15 minutes at 70C to help remove proteins non-specifically bound to the DBCO beads. After the incubation, samples were centrifuged for 5 minutes at 1000g and the resultant supernatant discarded. The DBCO agarose beads were resuspended in 0.5 mL 40 mM Iodoacetamide (Millipore Sigma, I1149-25G) and incubated light protected for 30 minutes to alkylate proteins. After the incubation, samples were centrifuged for 5 minutes at 1000g and the resultant supernatant discarded and the DBCO agarose beads were resuspended in 500 µL 0.8% SDS. The DBCO agarose beads were subjected to extensive washing to further remove non-specifically bound proteins. This was accomplished by washing each sample with 50 mL of 0.8% SDS, 8 M Urea, and 20% acetonitrile (Fisher Scientific, PI51101). The speed of washes was enhanced by performing them in Poly-Prep Chromatography Columns (Biorad, 7311550) connected to a vacuum manifold (Fisher Scientific, 7311550); approximately 7 mL of a wash was added to the column to resuspend the DBCO agarose beads, and then the vacuum applied to draw through the wash buffer, leaving DBCO agarose beads within the column. Following all washes, DBCO agarose beads were resuspended in 700 µL of 50mM HEPES (pH 8.0) (and immediately transferred to a 1.5 mL tube. DBCO beads were centrifuged for 5 minutes at 1000g. After centrifugation, the supernatant was completely removed and 200 µL 50 mM HEPES (pH 8.0) (Fisher Scientific, AAJ63002-AE) was added to the DBCO agarose beads. Next, 10 µL of a 0.1 ug/ µL Trypsin/Lys-C Mix (Fisher Scientific, V5073) was added to each sample. Proteins bound to the DBCO agarose beads were on-bead digested overnight at 37C on a thermomixer set to 1500-2000rpm. The next morning, approximately 16 hours after initiating on-bead digestion, samples were centrifuged for 10 minutes at 1000g. Supernatant containing digested peptides was transferred to a new tube and frozen at −80C until further processed. Peptide amounts were quantified with the Pierce Quantitative Peptide Assays & Standards kit (Thermo Fisher Scientific, 23290). Peptides destined for single shot LC-MS experiments were desalted using Nest Group Inc BioPureSPN Mini, PROTO 300 C18 columns (Fisher Scientific, NC1678001). The desalting involved conditioning the column with 200 µL methanol for five minutes followed by centrifugation at 25g until dry, washing the column twice with 200 µL 50% acetonitrile, 5% formic acid (Thermo Fisher Scientific, 28905) by centrifugation at 25g until dry, washing the column four times with 5% formic acid by centrifugation at 25g until dry, passing peptides through the column in 40 µL increments by centrifugation at 25g until dry, washing the column four times with 200 µL 5% formic acid by centrifugation at 25g until dry, and finally eluting the peptides two times with 100 µL 80% acetonitrile, 0.1% formic acid by centriguation at 25g. Following desalting, peptides were dried in a speed vac and then maintained at −80C before being run by LC-MS.

Peptides destined for tandem mass tagging (TMT) and pooling were dried in a speed vac and subsequently resuspended in 25 µL 100mM TEAB (pH 8.5) (Millipore-Sigma, T7408-100ML). TMTpro 18-plex reagents (Thermo Fisher Scientific, A52047) were reconstituted to 4 µg/uL in anhydrous acetonitrile (Millipore-Sigma, 271004-1L). A volume of TMT label was added to the peptide suspension to obtain a minimum of 10 µg TMT per 1 µg peptide and to maintain a TEAB to acetonitrile ratio of 5:2. Peptides were incubated with TMT labels for two hours with occasional vortexing. Labeling reactions were quenched by adding 2 µL 50% hydroxylamine (Thermo Fisher Scientific, B22202.AE) for 15 minutes with occasional vortexing. Equal volumes of TMT-labeled peptides were pooled and dried in a speed vac, after which peptides were desalted as described above for single shot LC-MS preparations.

### In-Solution Digestion of Proteins for LC-MS Preparation

In-solution digest was employed for experiments examining bulk brain proteome and aggregates from wildtype aged mice. Protein was chloroform-methanol precipitated by adding 600 µL methanol, 150 µL chloroform (Millipore-Sigma, C2432-500ML), and 400 µL water to 40 µL protein sample and centrifuging the mixture at 17,000g for 5 minutes. The upper phase was discarded and 650 µL methanol was added to the sample, vortexed, and centrifuged for 17,000g for 5 minutes. The supernatant was removed and the protein pellet dried for 10 minutes. The dried protein pellet was resuspended in 10 µL of 8 M urea, 0.1M Tris-HCL (pH 8.5). 2.25 µL 10 mM DTT was added to the sample and vortexed, followed by incubation on a thermomixer at 30C with shaking at 650rpm for 90 minutes. Next, 2.83 µL 50 mM IAA was added to the sample and vortexed, followed by light-protected incubation for 40 minutes. After the incubation, 90 µL 50 mM Tris (pH 8) was added to dilute the urea concentration. Lastly, Trypsin/Lys-C Mix was added to a mass to mass ratio of 1:50 and the samples were digested overnight at 30C with shaking at 650rpm on a thermomixer. Following digestion, peptides were desalted as described for preparation of BONCAT-labeled proteins.

### Mass Spectrometry Data Acquisition and Data Processing for Bruker timsTOF Pro Data

timsTOF was generally utilized for small-scale comparisons (8 or fewer samples to be directly compared) and/or when peptide amount was limited and high sensitivity was still desired. timsTOF was used to acquire the following data in this manuscript: brain region comparison data in Figure 1; neuronal aggregate data in Figure 4; neuron to microglia protein transfer data in Figure 5; CMV-Cre;BONCAT data from various tissues in the supplemental data; Samples are analyzed by TimsTOF Pro mass spectrometer (Bruker Daltonics) coupling with NanoElute system (Bruker Daltonics) with solvent A (0.1% formic acid in water) and solvent B (0.1% formic acid in Acetonitrile).

Dried peptides were reconstituted with solvent A and injected onto the analytical column: Aurora Ultimate CSI 25×75 C18 UHPLC column, by NanoElute system at 50 °C. The peptides were separated and eluted by the following gradient: 0 min 0% B, 0.5 min 5% B, 27min 30% B, 27.5min 95% B, 28min 95% B, 28.1min 2% B and 32 min 2% B at a flow rate of 300nL/min.

Eluted peptides were measured in DDA-PASEF mode using timsControl 3.0. The source parameters were 1400V for capillary voltage, 3.0l/min for dry gas and 180 °C for dry temperature using Captive Spray (Bruker Daltonics). The MS1 and MS2 spectra were captured from 100 to 1700 m/z in Data-Dependent Parallel Accumulation-Serial Fragmentation (PASEF) mode with 4 PASEF MS/MS frames in 1 complete frame. The ion mobility range (1/K0) was set to 0.85 to 1.30 Vs/cm2. The target intensity and intensity threshold were set to 20,000 and 2500 in MS2 scheduling with active exclusion activated and set to 0.4min. 27eV and 45eV of collision energies were allocated for 1/K0=0.85 Vs/cm2 and 1/K0=1.30 Vs/cm2 respectively.

Data captured was processed using Peaks Studio (Version10.6 built on 21st December 2020, Bioinformatics Solution Inc.) for sequence database search with the Swiss-Prot Mouse database. Mass error tolerance was set to 20ppm and 0.05Da for parent and fragment ions.

Carbamidomethylation of cysteine was set as a fixed modification. Protein N-term acetylation and methionine oxidation were set as variable modifications, with maximum of 3 variable PTM allowed per peptide. Estimate FDR with decoy-fusion is activated. Both FDR for peptides and proteins were set to 1% for filtering.

### Mass Spectrometry Data Acquisition and Data Processing for Thermo Eclipse Data

The Thermo Eclipse was used for any experiments utilizing TMT as this instrument is capable of running TMT samples. TMT, and by extension the Thermo Eclipse, were utilized for larger-scale comparisons to avoid any time-associated ‘drifts’ that would make comparisons between samples run on an instrument far apart in time less accurate. Additionally, TMT was utilized when quantitative precision and consistent peptide identification across replicates or samples to be compared were critical, such as for the protein degradation experiments in which the quantification of all time points of a single region and single age was instrumental to the overall success of the experiment. TMT labeling and the Thermo Eclipse were used for the following experiments: transgenic line comparisons in Figure 1; protein degradation experiments in Figures 2 and 3; AAV and transgenic mouse comparisons in the supplemental data; aged versus young and tauopathy versus aged matched wildtype in supplemental data.

Samples were analyzed by Easy-nLC 1200 coupled to the Thermo Scientific™ Orbitrap Eclipse™ Tribrid™ mass spectrometer with EasySpray Ion source and FAIMS Pro interface. Digested samples were reconstituted in 0.1% formic acid in water and were loaded to a trap column (Thermo Scientific™ Acclaim™ PepMap™ C18 column, 2 cm x 75 µm ID, 3 µm) and separated on Thermo Scientific™ Acclaim™ PepMap™ RSLC C18 column, 25 cm x 75 µm ID, 2 µm. Solvent A is 0.1% formic acid in water and solvent B is 80% acetonitrile in water with 0.1% formic acid. The gradient was ramped from 2% B to 40% B in 179 minutes at a flow rate of 300 nL/min. The column temperature was set at 50°C.

TMT labeled peptides were analyzed by data-dependent acquisition mode using Synchronous Precursor Selection (SPS) MS3 Real Time Search (RTS) approach. For full MS Scan, resolution was set at 60,000 and the mass range was set to 350-1500 m/z. Normalized AGC Target was set at 100% and the maximum injection time is 50ms. The most abundant multiply charged (Charge 2-7) parent ions were selected for CID MS2 in the ion trap. The CID collision energy was set at 35%. Real Time Search using Uniprot-Mus musculus database was performed. Carbamidomethyl on Cysteine (C) and TMTpro 16plex on lysine (K) and peptide terminal were set as static modifications. Oxidation on Methionine (M) was set as variable modifications. Up to 10 parent ions from MS2 will be selected by Synchronous Precursor Selection (SPS) for HCD MS3. MS3 spectra were acquired at 30,000 resolution (at m/z 200) in the Orbitrap MS with 55% normalized HCD collision energy. The cycle time was set at 2 seconds, and 3 experiments were run for different FAIMS Compensation Voltage (CV): −45V, −60V and −70V.

TMT data were processed using Thermo Scientific™ Proteome Discoverer™ software version 2.4. Spectra were searched against a UniProt Mus Musculus database using the SEQUEST® HT search engine. Maximum 2 missed cleavage sites was set for protein identification. Static modifications included carbamidomethylation (C) and TMTpro (K and peptide N-terminus). Variable modifications included oxidation (M) and acetylation (Protein N-terminus). Resulting peptide hits were filtered for maximum 1% FDR using the Percolator algorithm. The MS3 approach generated CID MS2 spectra for identification and HCD MS3 for quantitation. Precursor mass tolerance was set as 10ppm and fragment mass tolerance for CID MS 2 Spectra obtained by Ion Trap was set as 0.6 Da. The peak integration tolerance of reporter ions generated from SPS-MS3 was set to 20 ppm. For the MS2 methods, reporter ion quantification was performed on FTMS MS2 spectra and for identification, where they were searched with precursor mass tolerance of 10 ppm and fragment mass tolerance of 0.02 Da.

For the reporter ion quantification in all methods, no normalization and scaling were applied. The average reporter s/n threshold was set to 10. Correction for the isotopic impurity of reporter Quan values was applied.

### Mass Spectrometry Data Analysis: Basic Fold Change Over Background Analysis and Protein Identification

For experiments in which the goal was to simply quantify the number of different proteins labeled in BONCAT-labeled samples and/or identify proteins labeled with confidence over the background, non-normalized data was used as input. The reason for using non-normalized data is because the background samples were expected to have few proteins and low abundance relative to labeled samples, and this inherent difference should be preserved for analyses of labeled samples relative to background samples; normalizing would remove this inherent difference. The following steps were performed on the data: data was log_2_ transformed, proteins were filtered based on possessing valid values in a certain number of replicates in at least one group, missing values were replaced by imputation (width of 0.3 and downshift of 1.8), fold change was calculated for each protein between BONCAT-labeled replicates versus the respective background control replicates, and *p* values for these comparisons were derived from a two-tailed t-test. The number of replicates that were required to possess a valid value for a protein was dependent on the number of replicates used in the experiment, but in all cases required over 50% of the replicates in at least one group to possess a valid value for any given protein; the Camk2a-Cre;BONCAT benchmarking experiment required 3 of 4 replicates to have valid values in at least one group, the AAV-Camk2a;PheRS* experiment required 3 of 4 replicates to have valid values in at least one group; CMV-Cre;BONCAT experiments required 2 of 2 replicates to have valid values in at least one group, the neuronal aggregate experiment required 2 of 3 replicates to have valid values in at least one group, the neuronal protein transfer experiment required 2 of 3 replicates to have valid values in at least one group. *P* values less than 0.05 were considered significant in all analyses, with the fold-change cutoff varying by experiment and indicated in the respective figures. For analyses in which labeled protein was expected to be rare, as in the neuronal protein aggregate study and neuron to microglia protein transfer study, a less stringent fold change criteria was imposed (>0 over background controls). For analyses in which labeled protein was expected to be abundant, as in the benchmarking experiments in Figure 1, more stringent fold change criteria were imposed (> 2 fold change over background controls).

### Mass Spectrometry Data Analysis: Principal Component Analysis

For principal component analysis, individual data frames from each group being compared underwent filtering as described above in the basic fold change analysis with any proteins remaining after filtering in each group being retained as true hits. Subsequently, dataframes for each group containing the raw abundance values were merged with all proteins being retained regardless of whether they were shared or not among the three groups. The raw abundance values were log_2_ transformed and missing values, which were mostly proteins that were identified in one BONCAT line or region but not others, were replaced by imputation (width of 0.3 and downshift of 1.8). By implementing imputation, which is necessary for PCA, each sample possessed the same number and identity of proteins, not only the number shown in the Venn Diagram comparing the BONCAT mouse lines. These values were then used for principal component analysis in Perseus software.

### Mass Spectrometry Data Analysis: Comparing Protein Abundance Between Different Groups

In a few experiments, labeled protein fold changes between two or more experimental conditions which were also labeled were compared. The experiments include regional comparisons in Fig. 1 and aged versus young comparison in Extended Data Fig. 2. In these experiments, data frames from each group being compared underwent filtering as described above in the basic fold change analysis with any proteins remaining after filtering in each group being retained as true hits for further analysis. Data frames of true hits for each group were merged to keep all proteins identified among all groups. The raw abundance values for the retained proteins from all groups were log_2_ transformed and missing values, which were mostly proteins that were identified in one condition but not the other, were replaced by imputation (width of 0.3 and downshift of 1.8). The fold change for these values were calculated for each protein between conditions and *p* values for these comparisons were derived from a two-tailed t-test. In the case of the three-way regional comparison in Fig. 1, the resultant values were z-scored before visualizing in a heatmap.

### Mass Spectrometry Data Analysis: Protein Turnover Analysis

As described above, non-normalized data was used as input for protein turnover analyses with a similar rationale to preserve inherent differences in protein abundance between time points, with less protein being expected at each successive time point progressing into the chase period; normalization would risk losing these inherent differences. First, the log_2_ fold change in protein abundance between time point 1 replicates and wildtype background control replicates was calculated. Any proteins enriched > 1.5 in time point 1 over the background control were considered for further analysis and the other proteins were discarded as background proteins.

This fold change over background filtering was performed for each age and region combination separately. Time point 1 was used in this filtering approach rather than other time points because time point 1 represents the time point of maximal labeling and would give the fairest evaluation of background compared to later time points at which enriched proteins successively approach levels closer to that of wildtype controls and would lead to discarding more proteins likely unjustly. When making comparisons between different ages of a single region, proteins were further filtered to those that were commonly detected between ages and detected in all replicates with the only exception being the degradation kinetic trajectories shown in Fig. 2d. With this relatively stringent filtering approach, we ensure equally reliable half-life predictions and trajectory analysis without a need to assign uncertainty values due to varying drop-out rates.

For kinetic degradation trajectory analyses, the mean of each replicate of each time point in each age and region was calculated. Time point 1 was considered 100% protein remaining for each protein in each region and age and the other time point percentages were calculated by dividing the average abundance of the successive time point by the average abundance of time point 1.

Differences between all subsequent time points were calculated and only decreasing trajectories or trajectories with an up to 5% increase between two time points were retained. Because labeled protein should either decrease or remain stable during a pulse-chase experiment, we considered an increase above 5% between time points as caused by measurement noise, we elected to exclude proteins that exceeded this 5% threshold between any two consecutive time points. Of note, in a seminal study employing SILAC labeling *in vitro* to measure protein degradation^6^, a threshold of 130% protein remaining was used (see Fig. 1D in cited paper), which is much less stringent that which we apply. For this experiment, a few technical notes should be made: First, all replicates of all time points from one region and one age were labeled with tandem mass tags and combined into one plex to permit the most accurate quantitative analysis of protein degradation; Second, from the resulting data in each plex, for any given protein in one region and for one age, the amount of protein present/remaining at timepoint 1 was the maximum and considered 100% and the percent remaining in subsequent timepoints was the fraction of the average abundance of biological replicates at that timepoint divided by the average abundance of biological replicates of timepoint 1. To compare protein turnover between different regions and/or ages, the percentages of protein remaining at specific times were compared. Notably, by comparing the percent protein remaining between regions and ages rather than directly comparing raw abundance values, natural differences in protein synthesis and/or variability in protein labeling would not skew analyses and data interpretation. Trajectories were clustered using fuzzy c-means clustering. The optimal number of clusters was determined using minimal centroid distance. For comparison with the aged groups, matching proteins within that group were separated in the same cluster distribution. Integrals for each protein in each cluster were calculated based on the trajectories used for clustering. The integral for each protein was calculated by trapezoidal numerical integration using MATLAB trapz. Delta integral values were derived by calculating the difference between the integral value of the aged protein and the respective young protein, where average delta integral values were determined similarly with the addition of dividing by the number of different protein species analyzed (for example, the number of proteins in a certain cluster). The significance of the average delta integral values was determined by a one-way ANOVA with significant comparisons determined by a Tukey test.

For half-life estimations, we normalized the mean trajectory for each protein such that the first measurement at timepoint zero corresponds to the value of one. These normalized trajectories were fitted using the method of least squares, meaning that the objective function to minimize was defined as the sum of the square of the difference between the goal function and the data.

The functions describing the one-and two-level model were given as Λ_l_(t) = *exp*(-k*_A_*t) and Λ_2_(t) = G(t) - G(7+ t_p_)/G(0) - G(t_p_) with G(t) = (k_AB_ (k_AB_ + k_A_) exp(-k_B_ t) + k_B_ (k_A_ - k_B_) exp(-(k_AB_ + k_A_)t)).

The parameters minimizing the objective function were then found using Matlab fmincon. For the one-level model, we calculated the half-life directly from the decay rate k_A_. The half-life for the two-level model was found by linear interpolation. After calculating the AIC AIC = n log(RSS/n) + 2k) for both models, we chose the model with the lower AIC and its corresponding half-life. These modeling approaches were derived from previous publications that studied protein degradation by pulse-chase methodology and subsequently estimated half-life^6,7^. It is important to note that as shown in the extended data, the modeling approach was in good correlation with direct interpolation of half-lives from degradation trajectories. Because of the good correlation and the benefit of being able to estimate the half-life of proteins that did not reach or go below 50% remaining in our data.

### Extracting Protein Clusters from Heatmap

To extract the proteins that defined the clusters, and by extension brain regions, in the z-scored heatmap comparing the striatum, hippocampus, and motor cortex, hierarchical clustering information was extracted from the heatmap. First, the command “pl$tree_row” was used to extract hierarchical clustering information from the heatmap for the protein clusters. Next, the hierarchical protein tree was divided into three clusters by the “cutree” function, with three clusters chosen on the visual distinction of three gene clusters in the heatmap. Lastly, protein IDs were extracted by the command “which(lbl=x)”, where “x” represents the cluster number.

### Protein Feature Analysis

Protein features were extracted from a comprehensive table of proteins and protein features from a previous publication^8^ and matched to the proteins of interest in this manuscript. Comparisons of protein features were made between the groups of interest as reported in the main text with statistical analyses being performed either by t-test or one-way ANOVA with a Tukey Test.

### Gene Ontology Analysis

Gene Ontology analyses were performed using the ShinyGO web application (http://bioinformatics.sdstate.edu/go/)^9^. Protein lists converted to standard gene symbols were uploaded to the application as input. Default parameters were used to run the analysis. For experiments related to protein aggregates and protein transfer, all proteins identified among all neuronal BONCAT models were used as a background gene list. The output, visuals and tables including enriched terms, enrichment FDR, number of genes in the pathway, and fold enrichment, were filtered by statistical significance (FDR < 0.05) and reported in this manuscript.

### Cell Type Annotation of Proteins

To annotate cell types, we utilized the ClusterMole R package (version 1.1). This package leverages a curated database of cell type marker genes to assign cell type probabilities to differentially expressed genes (DEGs). Specifically, ClusterMole compares the set of up- and down-regulated genes identified to known cell type marker signatures. A hypergeometric test is employed to calculate p-values for overrepresentation of cell type signatures within the DEG sets. When no cell type was annotated to a particular protein, it was considered non-cell type specific.

### Overlap with MAGMA-H

Neurodegenerative and neurodevelopmental risk genes were derived from a Hi-C multimarker analysis of genomic annotation (MAGMA) study^10^. By analyzing gene regulatory relationships in the disease-relevant tissue, this study identified neurobiologically relevant target genes, improving upon existing MAGMA studies. Lists of adult brain risk genes and summary statistics were downloaded from the studies GitHub repository at https://github.com/thewonlab/H-MAGMA. Genes with reported *p* values < 0.05 were considered risk genes.

### Signal Peptide Analysis

Signal peptide prediction was performed by querying protein sequences in SignalP, a server that predicts the presence of signal peptides and the location of their cleavage sites in proteins from Archaea, Gram-positive Bacteria, Gram-negative Bacteria, and Eukarya^11^. Individual protein sequences were retrieved from Uniprot and entered one by one into the SignalP browser search (https://services.healthtech.dtu.dk/services/SignalP-6.0/). We considered protein sequences with signal peptide scores > 0.1 to contain a bonafide signal peptide sequence and to be considered secreted. Proteins with signal peptide scores < 0.1 but > 0.02 were considered to contain a likely signal peptide sequence. Proteins with signal peptide scores < 0.02 were considered to unlikely contain a bonafide signal peptide sequence and thus considered unlikely to be secreted. Details of SignalP analysis can be found in the original publication^11^.

### ExoCarta Analysis

Classification of proteins as exosome cargo was performed by querying proteins on ExoCarta, a manually curated web-based compendium of exosomal proteins, RNAs and lipids^12^. Individual gene symbols or protein names were entered one by one into the ExoCarta browser query search (http://exocarta.org/query.html). If the search resulted in any mammalian hit, it was considered a potential exosome cargo. Details of ExoCarta analysis can be found in the original publication^12^.

### SynGO Analysis

An in-depth analysis of synaptic ontologies of a protein list was performed by using SynGO, an evidence-based, expert-curated resource for synapse function and gene enrichment studies^13^.

Gene lists were input to the SynGO browser (https://www.syngoportal.org) and default analysis parameters were applied. Visualizations of enrichment analysis on SynGO Cellular Components and Biological Processes were exported from the SynGo browser. Details of SynGO analysis can be found in the original publication^13^.

### Uniprot ID to Gene Symbol Conversion

Uniprot IDs were converted to gene symbols using Uniprot’s Retrieve/ID mapping web tool (https://www.uniprot.org/id-mapping). In cases in which multiple gene symbols were returned for a single Uniprot ID, the entry name – the unique gene symbol identifier associated with the Uniprot ID - was used for most in-text references and visualizations. All gene symbols associated with a single Uniprot ID are listed within the extended data tables with the entry name listed first in the list.

### Mining of Pre-print data

Analysis of mouse and human microglia proteomes was performed on processed LC-MS data from a pre-print^14^. The reported copy number of the four replicates of freshly isolated 3.5 month-old male mouse microglia were averaged, and any protein with an average copy number >0 was considered as detected. This same analysis was performed on the five replicates of freshly isolated human microglia, derived from females aged 6 years, 22 years, 22 years, 45 years, and 61 years.

### Hypergeometric Test for Protein Overlap and Enrichment

Hypergeometric p values and related enrichment values were calculated by the scipy.stats package in Python 3.9. The background protein list/number used for these tests was 3787, the number of different BONCAT-labeled neuronal proteins we could maximally detect in the Camk2a;PheRS* model as shown in Fig. 1. All other numeric inputs were derived from the Venn Diagram displayed in Fig. 5k.

### Statistics

Details of statistical methods are described in relevant subsections above and/or indicated in figure legends. All t-tests were two-tailed. ANOVAs were ordinary one-way ANOVAs.

### Data Visualizations

Except if stated otherwise in the above methods, data visualizations were performed in R studio (Posit Software, Boston, MA, USA), GraphPad Prism (GraphPad Software), or Adobe Illustrator (Adobe, CA, USA) with aesthetic enhancements performed in Adobe Illustrator. Renderings of mice in Fig. 2a and Extended Data Fig. 2h, 2k and tau tangles in Extended Data Fig 2k were derived from Biorender.com.

### Data Availability

The raw mass spectrometry proteomics data have been deposited to the ProteomeXchange Consortium via the PRIDE partner repository. Pre-publication, data is accessible with a token only to reviewers. Once published, no token will be required, and the data will be freely accessible.

For datasets related to comparing BONCAT models in the context of a Camk2aCre driver, project accession PXD057020. File name annotations indicate the BONCAT line (MetRS*, PheRS*, or TyrRS*) examined within the dataset, which consists of TMT-plexed samples containing BONCAT-labeled samples and respective wildtype background controls. For the MetRS* dataset, samples 1676-79 are background controls and 1538-41 are MetRS* labeled samples. For the PheRS* dataset, samples 910-12 are background controls and 1584-90 are PheRS* labeled samples. For the TyrRS* dataset, samples 1701-04 are background controls and 1482-83 and 1471-72 are TyrRS* labeled samples.

For CMVCre;BONCAT datasets, project accession PXD056569. File name annotations are as follows: WT in name indicates sample is a background control (no BONCAT-labeleing); 4 digit code to start file name indicates sample is a BONCAT-labeled sample. Abbrievation following WT or 4 digit code indicates tissue type (B = brain, Liv = liver, H = heart, I = intestine, Lug = lung, M = muscle).

For datasets related to BONCAT-labeled neuronal protein differential expression by region, project accession PXD057261. File name annotations are as follows: Number indicates mouse ID; FC, Hipp, or ST annotation refers to brain region of sample (FC = Motor Cortex, ST = Striatum, Hipp = Hippocampus); TG annotation indicates the mouse was a transgenic BONCAT mouse in which protein labeling occurred; WT annotation indicated the mouse was a wildtype mouse in which BONCAT labeling could not occur and is thus a background control.

For dataset related to comparing BONCAT labeling in Camk2aCre;PheRS* knock-in mice to mice transduced with AAV-Camk2a;PheRS*, project accession PXD057456. Samples 1584-86 are BONCAT-labeled Camk2aCre;PheRS* knock-in mice; samples 1947-49 are BONCAT-labeled AAV-Camk2a;PheRS* transduced mice; samples 1400-02 are non-BONCAT labeled background controls.

For dataset related to comparing aged and young proteomes from mice transduced with AAV-Camk2a;PheRS*, project accession PXD057488. File name annotations are as follows: Y or A annotation in file name indicates whether sample was from a young (Y) or aged (A) mouse; TP1 in file name indicates sample was BONCAT labeled; BG annotation in file name indicates sample was a non-BONCAT labeled background control.

For datasets related to protein degradation among brain regions and tissues, project accession PXD056701. File name annotations are as follows: Y, M, or A annotation in file name indicates whether the plex consists of young, middle-aged, or aged samples, respectively. Number preceding Y, M, or A represents the brain region in the plex (3 = sensory cortex, 4 = visual cortex, 6 = hippocampus, 8 = hypothalamus).

For datasets related to BONCAT-labeled neuronal proteins in aged aggregates, project accession PXD056972. File name annotations are as follows: BG annotation in file name indicates the sample (n = 3) was a background controls; BON annotation in file name indicates the sample (n = 4) was derived from a BONCAT-labeled model.

For datasets related to label-free aged brain aggregates, project accession PXD057455. All files are replicates of label-free aggregates from the aged brain.

For datasets related to BONCAT-labeled neuronal proteins in microglia, project accession PXD056974. File name annotations are as follows: BG annotation in file name indicates the sample (n = 4) was a background controls; no BG annotation in file name indicates the sample (n = 3) was derived from a BONCAT-labeled model.

### Code Availability

Prepublication, unique codes generated and/or modified and used in this study to analyze the data are available from GitHub with a token only supplied to reviewers. Once published, no token will be required, and the repository will be freely accessible.

---

**Extended Data Figure 1:**
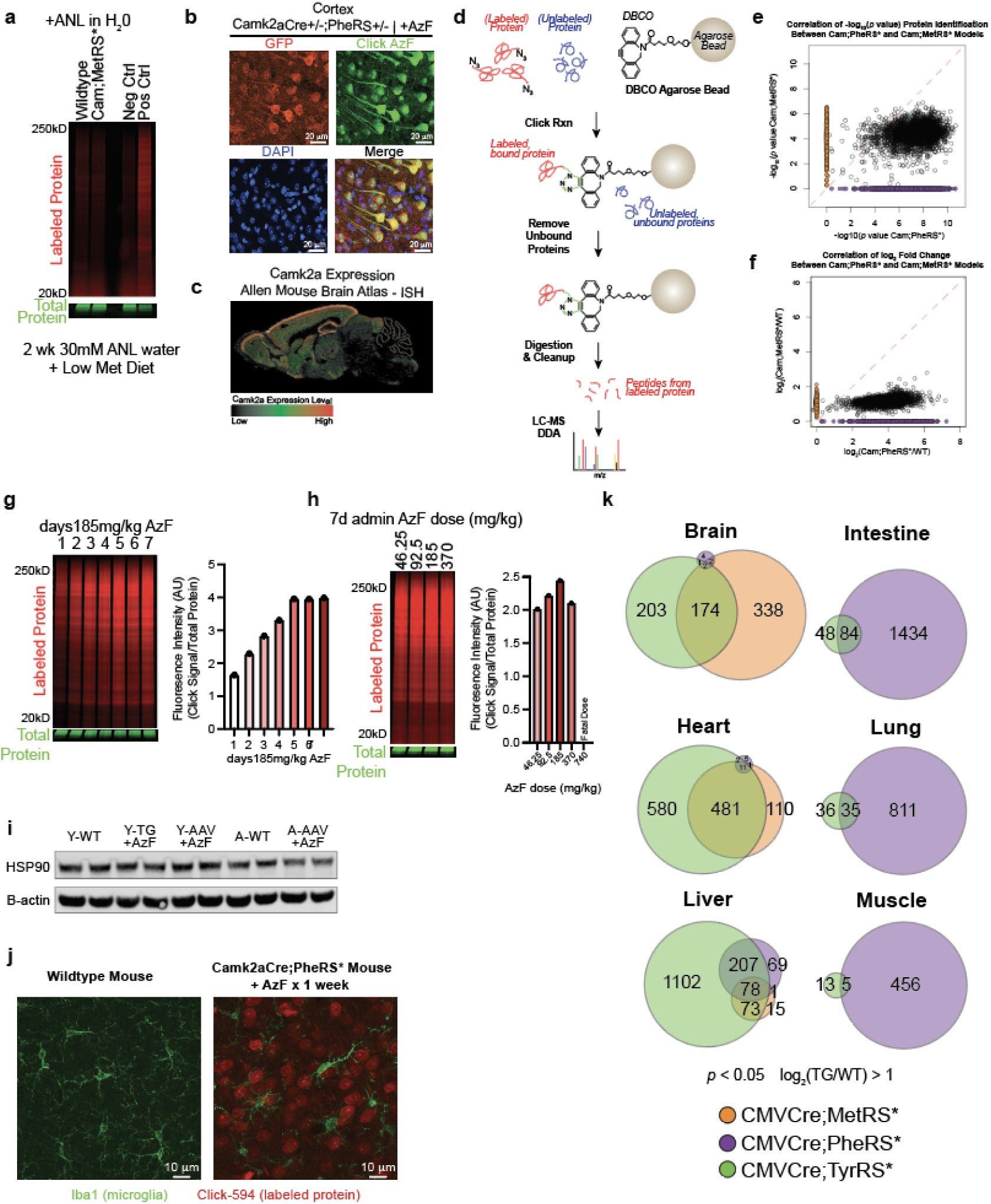
Evaluation of Nascent Proteome Labeling by Different BONCAT Mouse Lines. a. In-gel fluorescence image of Alexa 647-clicked and thus BONCAT labeled proteins of total brain lysates derived from young Camk2aCre;MetRS* model and its respective background control. These mice were provided 30mM ANL in water for 2 weeks while on a low methionine diet as reported in a protocol using the same BONCAT mouse line by the originators of the MetRS mouseline. b. Fluorescence images of Alexa 488-clicked and thus BONCAT labeled proteins in cortical tissue sections from young Camk2aCre;PheRS* model. Tissues are co-stained with anti-GFP, which should be co-expressed in all cells expressing PheRS*, and DAPI. c. *In situ* heatmap of Camk2a mRNA expression from the Allen Brain Atlas. d. Schematic of methodology used to enrich BONCAT-labeled proteins from total lysates for LC-MS. e. Scatter plot showing correlation of -log_10_ *p* values of proteins identified in the Camk2aCre;PheRS* model versus the Camk2aCre;MetRS* model. f. Scatter plot showing correlation of -log_2_ fold change (BONCAT/background control) of proteins identified in the Camk2aCre;PheRS* model versus the Camk2aCre;MetRS* model. g. In-gel fluorescence image of Alexa 647-clicked and thus BONCAT labeled proteins of total brain lysates derived young Camk2aCre;PheRS* mice provided 185mg/kg of azido-phenylalanine (AzF) for a varying number of days (left) and associated quantification of fluorescence intensity of clicked-protein normalized to total protein with the different number of days provided AzF (right). h. In-gel fluorescence image of Alexa 647-clicked and thus BONCAT labeled proteins of total brain lysates derived from young Camk2aCre;PheRS* mice provided varying doses of azido-phenylalanine (AzF) for one week (left) and associated quantification of fluorescence intensity of clicked-protein normalized to total protein with the different doses of AzF provided (right). i. Western blot image of HSP90 and loading control beta-actin on whole brain lysates derived from various BONCAT-labeled models and ages and respective non-labeled controls to show whether BONCAT-labeling induces an HSP90-mediated heat shock response. j. Fluorescence images for microglia (Iba1, green) staining in cortical tissue sections of wildtype, non-BONCAT labeled mice (left) and Camk2aCre;PheRS* BONCAT-labeled mice (right) to show whether BONCAT-labeling induces microgliosis as evaluated by cellular morphology. k. Venn Diagrams showing the overlap and exclusivity of proteins labeled by the CMVCre;MetRS*, CMVCre;PheRS*, and CMVCre;TyrRS* models in the indicated tissues. n = 2 biological replicates per group. Only proteins with a log_2_ fold change (BONCAT/background control) > 1 and *p* value < 0.05 were considered in this analysis.

**Extended Data Figure 2:**
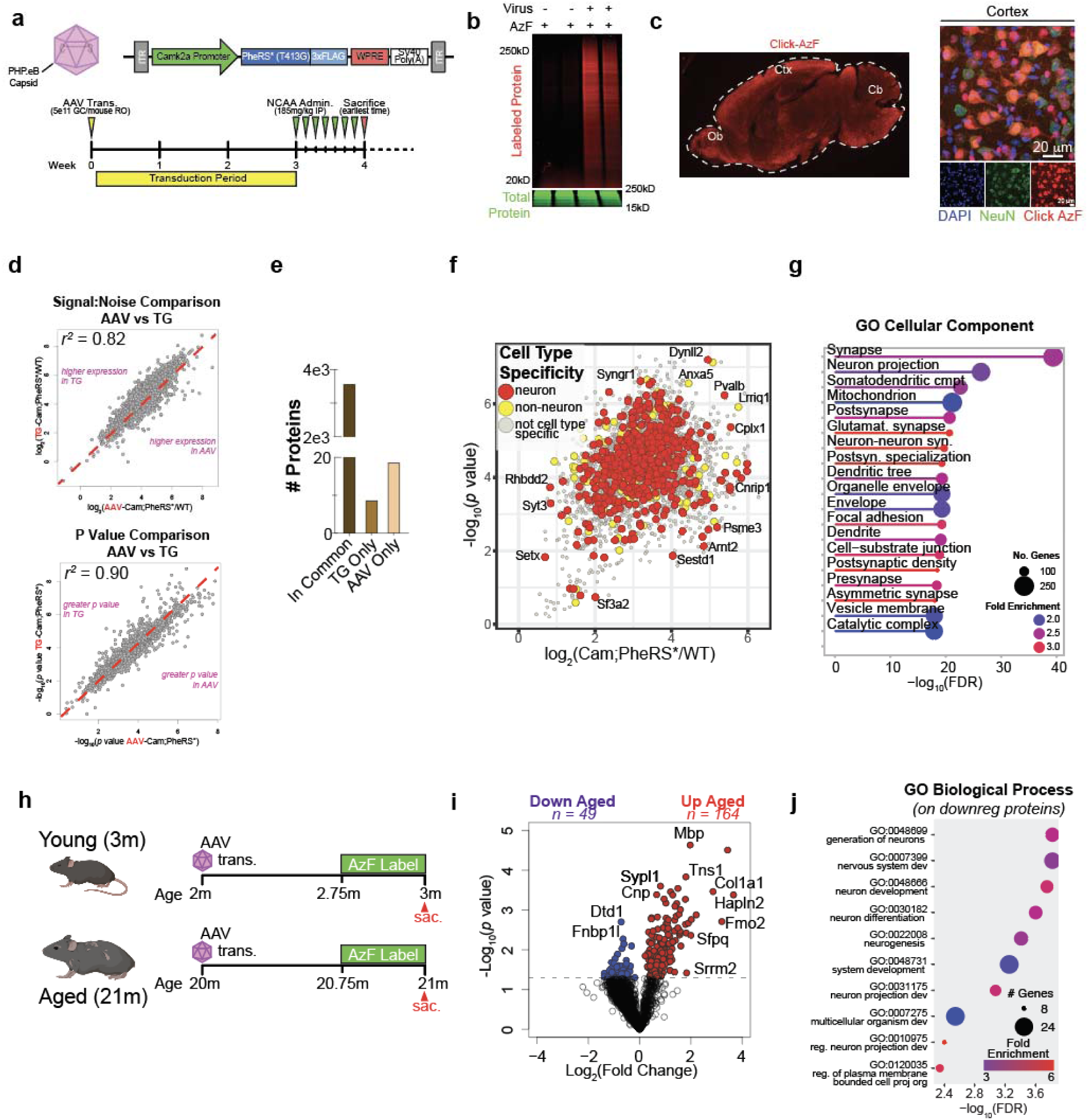
Evaluation of Nascent Proteome Labeling by AAV-Based Delivery of PheRS*. a. Schematic of AAV-expression construct for Camk2a promoter-driven expression of PheRS* (top) and experimental timeline of transduction and proteome labeling (bottom). b. In-gel fluorescence image of Alexa 647-clicked and thus BONCAT labeled proteins of total brain lysates derived from young (3 months) AAV-Camk2a;PheRS* transduced-mice and the respective background controls. c. Fluorescence images of Alexa 594-clicked and thus BONCAT labeled proteins in brain tissue sections from young AAV-Camk2a;PheRS* transduced-mice. The image on the right shows co-staining for neurons (NeuN, green) to show overlap between click signal and neurons as would be expected from this model. d. Scatter plots showing the correlation of -log_2_ fold change (BONCAT/background control) (top) and -log_10_ *p* values (bottom) of proteins identified in young Camk2aCre;PheRS* transgenic mouse model compared to that of the young AAV-Camk2a;PheRS* model. Only proteins commonly detected with a log_2_ fold change over respective wildtype background controls and *p* value < 0.05 were plotted. e. Bar chart showing the number of proteins identified by LC-MS commonly and exclusively in the young Camk2aCre;PheRS* transgenic mouse model and young AAV-Camk2a;PheRS* model. Only proteins with a log_2_ fold change over respective wildtype background controls and *p* value < 0.05 were used. f. Color-coded volcano plot showing the enrichment of BONCAT-labeled proteins identified by LC-MS in the young AAV-Camk2a;PheRS* model relative to the wildtype background control. Proteins are color-coded by cell type enrichment. g. Gene Ontology Cellular Component analysis on BONCAT labeled proteins in the young AAV-Camk2a;PheRS* BONCAT model. Proteins used in the analysis had a log_2_ fold change > 1 over the respective background control with a *p* value < 0.05. h. Schematic of AAV-Camk2a;PheRS* transduction and labeling in an experiment to compare nascent neuronal proteomes of young (3m) and aged mice (21m). n = 4 biological replicates per BONCAT-labeled sample per experimental group, n = 3 biological replicates per background control sample per experimental group. i. Volcano plot of neuronal proteins differentially expressed between young and aged mice. j. Gene Ontology Biological Process analysis on neuronal proteins downregulated in aged mice relative to young mice. Downregulated proteins were those with a log_2_ fold change < 0 and *p* value < 0.05, color-coded in blue in (i).

**Extended Data Figure 3:**
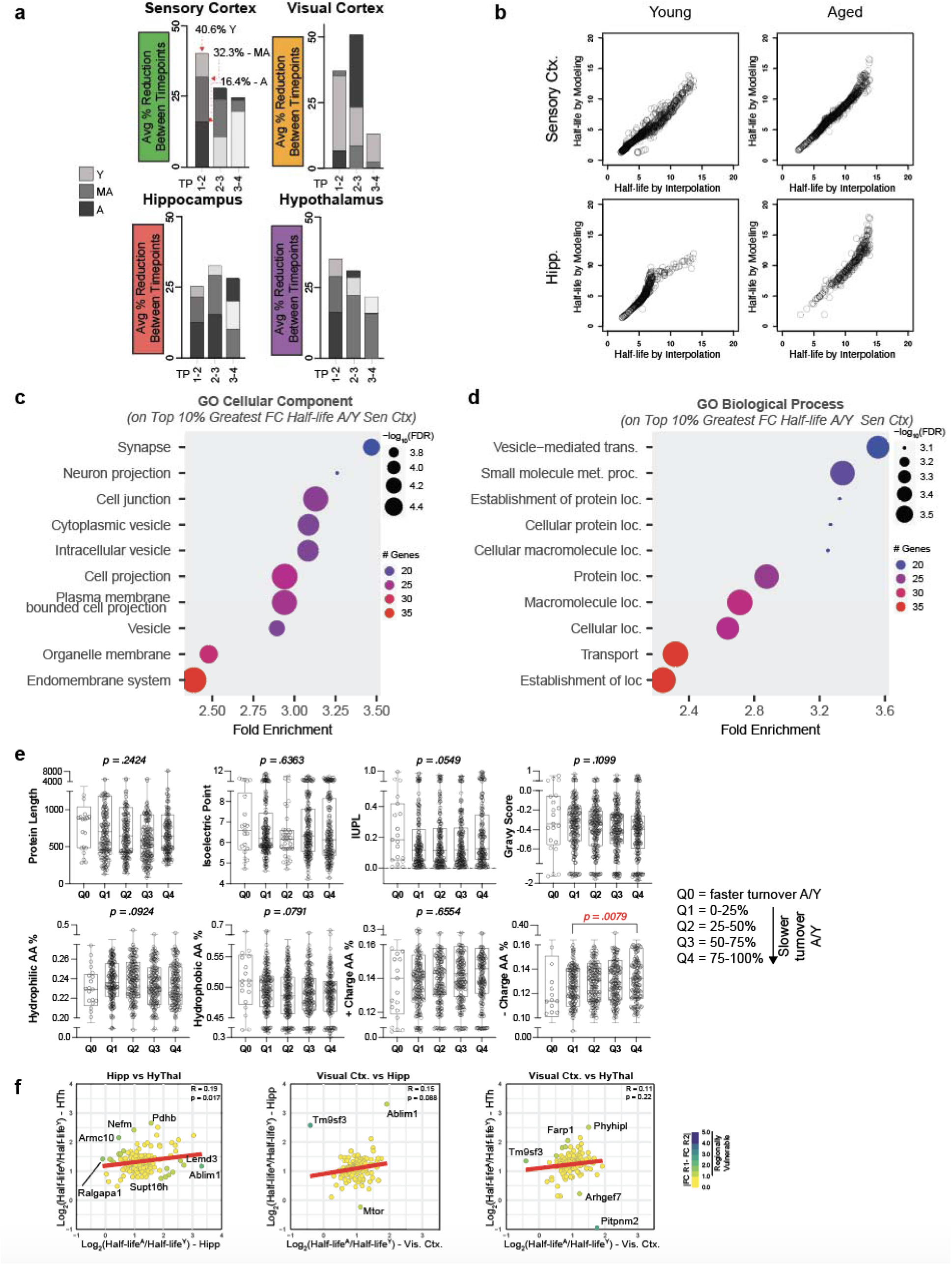
Analysis of Neuronal Protein Half-life with Aging. a. Bar charts showing the average percent reduction of neuronal protein abundance between consecutive time points for each age in each region analyzed. Bars should be interpreted as follows: wherever the top of the bar reaches along the y axis is the percent represented by that bar. b. Scatter plots showing the correlation of neuronal protein half-life in days estimated by modeling versus directly interpolated from the kinetic degradation plots. Only proteins that reached or surpassed 50% remaining are plotted because direct interpolation can only measure such proteins. c. Gene Ontology Cellular Component analysis of neuronal proteins from the sensory cortex within the top 10% greatest fold change (reduced degradation) from young to aged. d. Gene Ontology Biological Processes analysis of neuronal proteins from the sensory cortex within the top 10% greatest fold change (reduced degradation) from young to aged. e. Box and whisker plots comparing properties of neuronal protein from the sensory cortex within different quartiles of half-life fold change with aging. *P* values derived from a one-way ANOVA with significant comparisons identified by a Tukey test. f. Scatter plots comparing the log_2_ fold change of estimated protein half-lives (young to aged) between proteins commonly detected between the indicated regions. Each dot represents one protein with the color coding representing the absolute value of the difference between log_2_ fold changes of protein half-life between the indicated regions. Proteins with an absolute value difference >1 were considered regionally vulnerable.

**Extended Data Figure 4:**
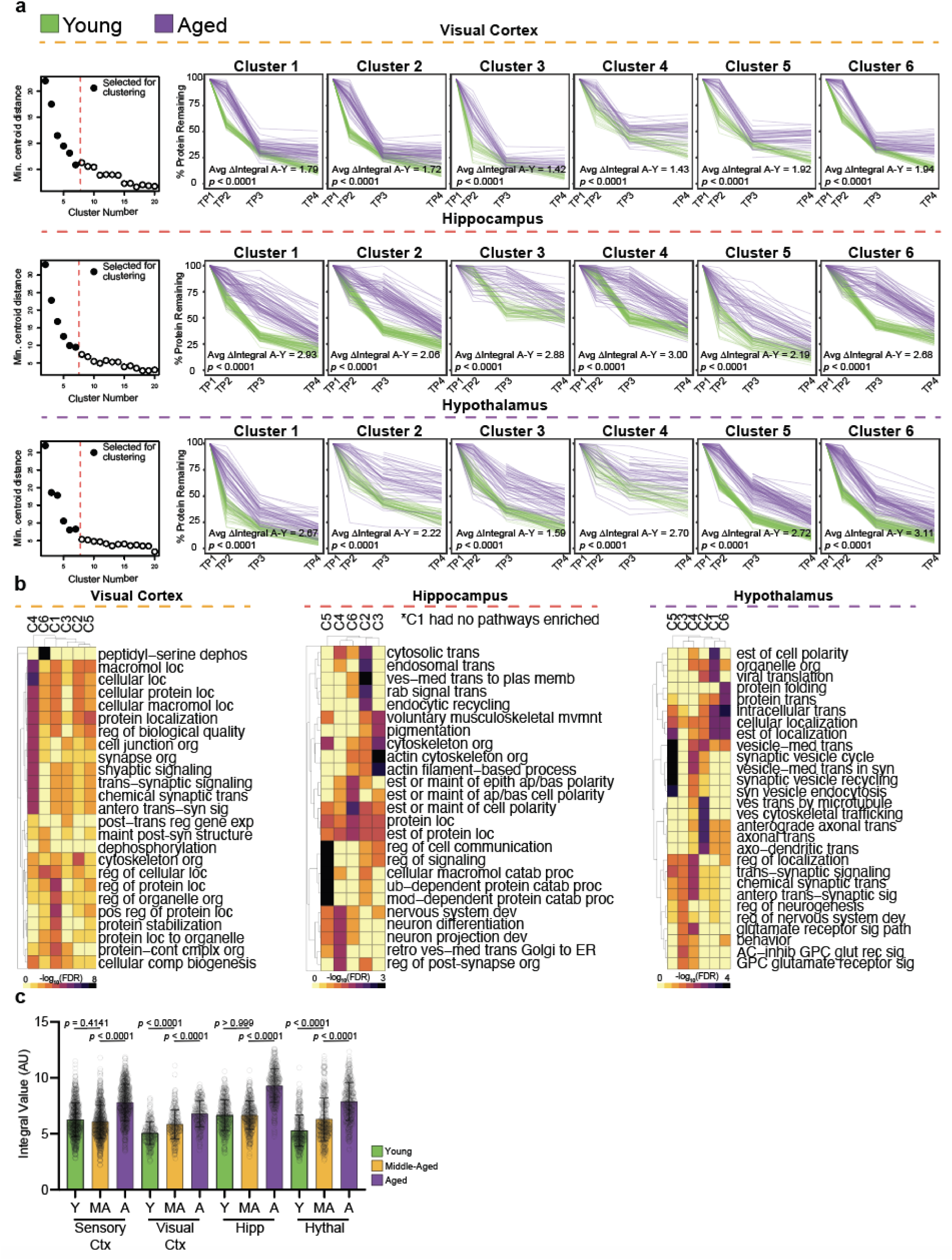
Clustering of Protein Kinetic Degradation Trajectories. a. Elbow plots of cluster number by minimum centroid distance used to determine cluster number for subsequent clustering analyses of young kinetic degradation trajectories with cluster cutoff indicated by a red dotted line (left) and associated clustering and overlap of young and aged kinetic degradation trajectories of the indicated brain regions. Protein membership in the aged clusters was determined by the clustering of young samples to serve as a baseline. The average delta integral, calculated by averaging the difference of the integral values of each aged and young protein within the cluster, is reported on each plot. The *p* value was determined by a one-way ANOVA with significant comparisons identified by a Tukey test (right). b. Heatmap of the top 5 most significant Gene Ontology Biological Processes identified for each cluster in the young visual cortex (left), hippocampus (middle), and hypothalamus (right). Heatmap colors represent -log_10_ of the FDR for each pathway. c. Bar plot comparing the integral values of young, middle-aged, and aged proteins on a per-region basis. *P* value determined by a one-way ANOVA with significant comparisons identified by a Tukey test.

**Extended Data Figure 5:**
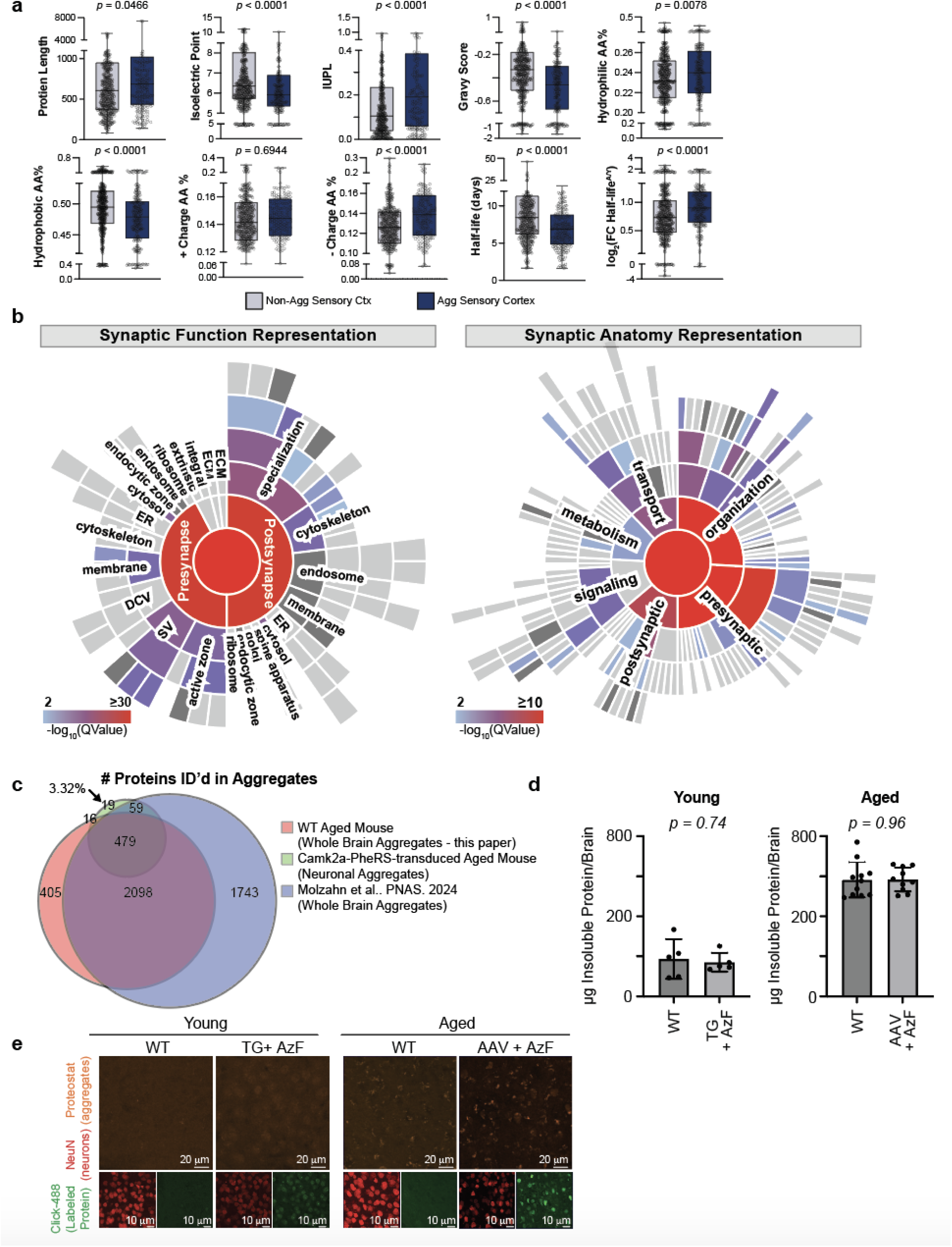
Analysis of Aged Neuronal Protein Aggregates. a. Box and whisker plots comparing properties of neuronal protein from the sensory cortex identified in aged neuronal aggregates compared to those not identified in aged neuronal aggregates. *P* values derived from a one-way ANOVA with significant comparisons identified by a Tukey test. b. Sunburst plots showing synaptic functional representation (left) and synaptic anatomical representation (right) of neuronal proteins identified in aged protein aggregates. c. Venn Diagram showing the overlap of neuronal proteins identified in aged protein aggregates by BONCAT methodology with proteins identified in aged protein aggregates without labeling methodology by us and an independent publication by Molzahn *et al*. d. Bar charts comparing mass of insoluble protein/protein aggregates between young Camk2aCre;PheRS* mice provided azido-phenylalanine (AzF) and young wildtype mice not provided AzF (left) and aged AAV-Camk2a;PheRS* transduced mice provided azido-phenylalanine (AzF) and aged wildtype mice not provided AzF (right). e. Fluorescence images comparing protein aggregate (Proteostat, orange) between young Camk2aCre;PheRS* mice provided azido-phenylalanine (AzF) and young wildtype mice not provided AzF (left) and aged AAV-Camk2a;PheRS* transduced mice provided azido-phenylalanine (AzF) and aged wildtype mice not provided AzF (right).

**Extended Data Figure 6:**
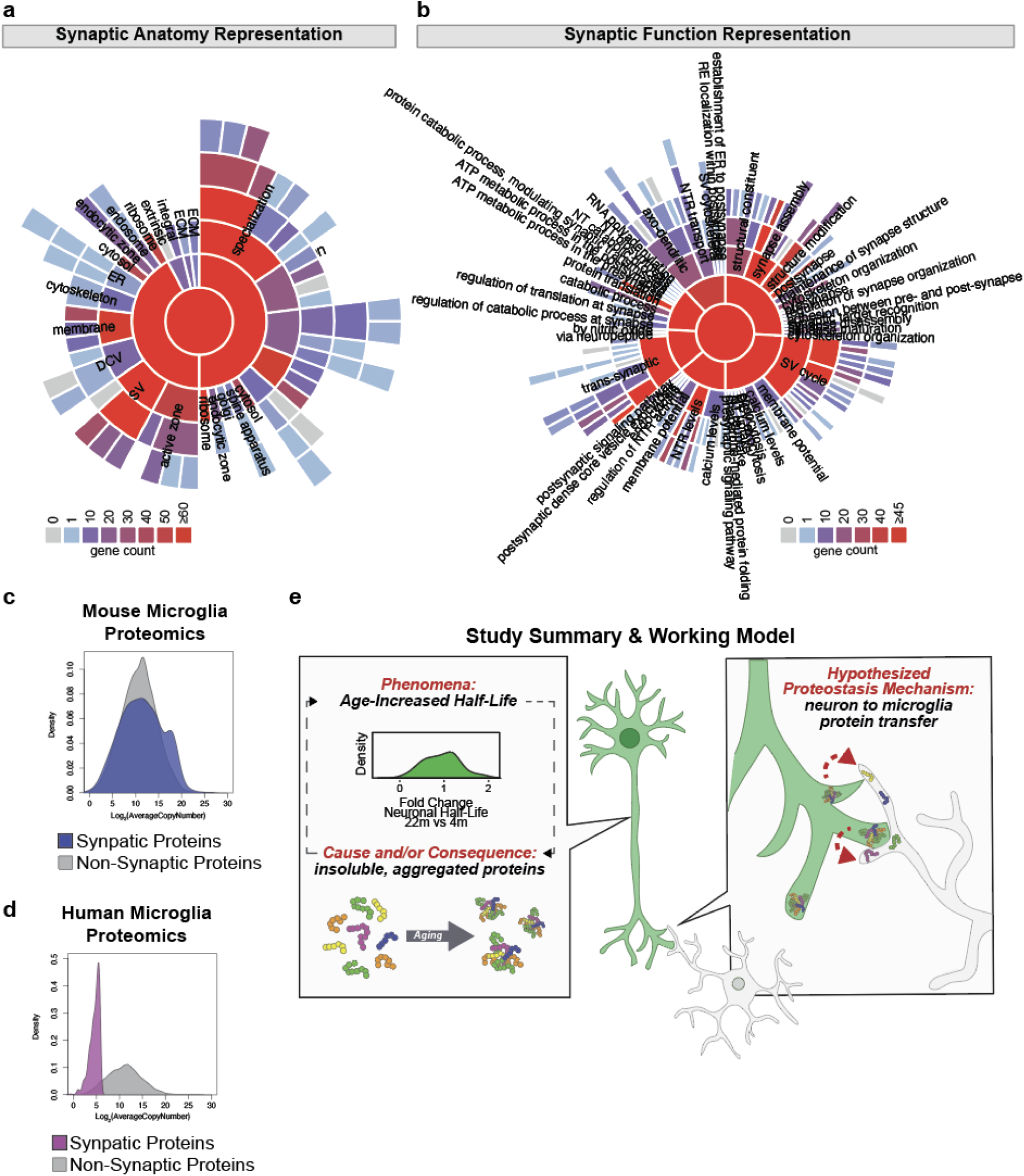
Analysis of Synaptic Proteins Found in Brain Macrophages. a. Sunburst plots showing synaptic anatomical representation of neuronal proteins identified in microglia. b. Sunburst plots showing synaptic functional representation of neuronal proteins identified in microglia. c. Density plot comparing the abundance of synaptic proteins and non-synaptic proteins in mouse microglia measured by LC-MS from Lloyd *et al*. d. Density plot comparing the abundance of synaptic proteins and non-synaptic proteins in human microglia measured by LC-MS from Lloyd *et al*. e. Schematic of study summary and working model.

